# Pioneer transcription factors coordinate active and repressive gene expression states to regulate cell fate

**DOI:** 10.1101/2022.12.29.522251

**Authors:** Satoshi Matsui, Marissa Granitto, Morgan Buckley, Joseph Shiley, William Zacharias, Christopher Mayhew, Hee-Woong Lim, Makiko Iwafuchi

## Abstract

Pioneer transcription factors (TFs) regulate cell fate by establishing transcriptionally primed and active states. However, cell fate control requires the coordination of both lineage-specific gene activation and repression of alternative lineage programs, a process that is poorly understood. Here, we demonstrate that the pioneer TF Forkhead box A (FOXA), required for endoderm lineage commitment, coordinates with the PR domain zinc finger 1 (PRDM1) TF to recruit Polycomb repressive complexes, which establish bivalent enhancers and repress alternative lineage programs. Similarly, the pioneer TF OCT4 coordinates with PRDM14 to repress cell differentiation programs in pluripotent stem cells, suggesting this is a common feature of pioneer TFs. We propose that pioneer and PRDM TFs coordinate recruitment of Polycomb complexes to safeguard cell fate.

**One-Sentence Summary:** Pioneer and PRDM transcription factors repress alternative lineage programs.

## Main Text

Pioneer transcription factors (TFs) prime and initiate gene regulatory programs, establishing developmental competence to differentiate into specific developmental lineages (*1*). For example, during endoderm differentiation, the pioneer TF FOXA locally open chromatin and recruits a Trithorax complex to deposit H3K4me1 active marks at endoderm-specific enhancers for priming transcription of endoderm genes (*2*–*5*). More broadly, the pioneering activity of Oct4, Sox2, and Klf4, along with c-Myc, enables somatic cells to be reprogrammed into pluripotent stem cells (*6, 7*). Repressive Polycomb chromatin domains, which help prevent ectopic gene expression, are established by Polycomb repressive complex 1 (PRC1), which ubiquitinylates K119 on histone H2A (H2AK119ub1), and PRC2, which methylates K27 on histone H3 (H3K27me3) (*8*–*11*). Polycomb chromatin domains exist in a bivalent state, with active histone modifications (H3K4me1/2/3) forming a transcriptionally poised state, primed for lineage-specific transcription but silenced until activated by the appropriate signals(*12*–*16*). However, how these bivalent domains are established during cell differentiation is unclear.

The FOXA family of TFs are encoded by three genes, *FOXA1, FOXA2*, and *FOXA3*, which are redundantly required for liver development from the embryonic endoderm (*17, 18*). To assess the impact of losing all FOXA factors on gene regulation in endoderm development, we established a doxycycline (dox) inducible CRISPR interference (CRISPRi) system in human pluripotent stem cells (hPSCs) to simultaneously knock down all three *FOXA* genes (**Fig. S1A**) (*19*) and used an endoderm differentiation protocol to generate homogenous cell populations of mesendoderm (ME), definitive endoderm (DE), foregut (FG), and liver bud progenitor (LBP) from hPSCs (**Fig. S1B**) (*20*). Dox treatment of *FOXA1/A2/A3-*CRISPRi (*FOXA-*TKD) hPSCs at the initiation of endoderm differentiation resulted in > 95% gene knock-down efficiency for all *FOXA* genes at the foregut stage, a period at which all *FOXA* genes are robustly expressed in dox-untreated control conditions (**Fig. 1A, S1B, S1C**).

**Fig. 1.**
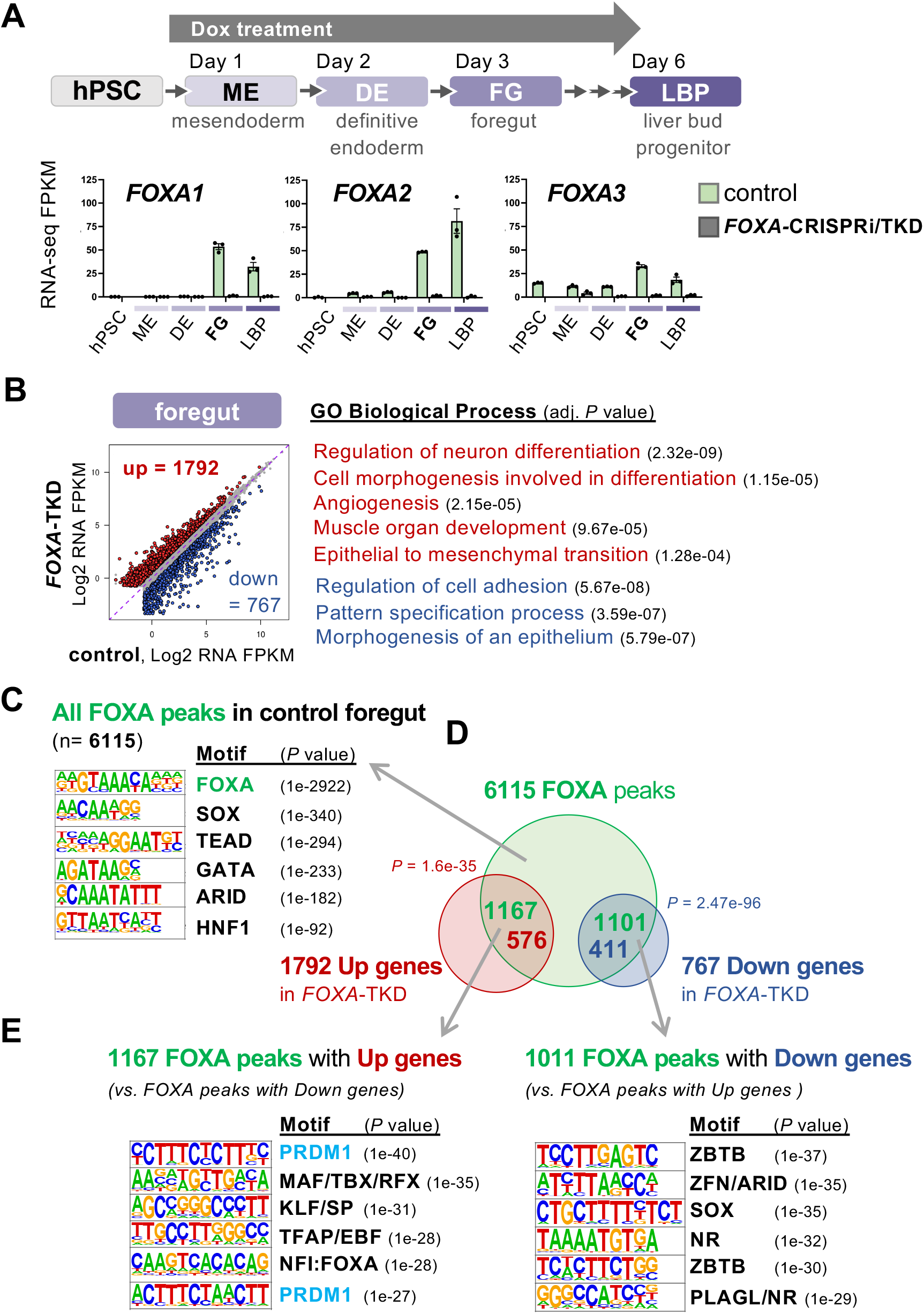
Pioneer TF FOXA prevents alternative-lineages and precocious gene expression during human foregut endoderm development. **(A)** Validation of *FOXA*-CRISPRi/TKD knockdown efficiency by RNA-seq during hPSC to endoderm differentiation (n=3 replicates from 3 independent *FOXA*-CRISPRi clones, Means ± SEM). **(B)** Differential gene expression analysis of RNA-seq comparing FOXA-TKO versus control and representative Gene Ontology (GO) term for upregulated (red) and downregulated (blue) genes in *FOXA*-TKD foregut (n=3 replicates; FDR < 0.05, FC > 1.5). **(C)** Top motifs enriched at combined FOXA1 and FOXA2 CUT&RUN peaks (FOXA peaks) in control foregut cells by HOMER (n=2 replicates from 2 independent dox-untreated *FOXA*-CRISPRi clones). **(D)** Intersections of FOXA peaks with differentially expressed genes (DEGs) in *FOXA*-TKD foregut by GREAT. *P* value of the overlaps by hypergeometric test. **(E)** Top motifs enriched at FOXA peaks associated with upregulated genes against a background of FOXA peaks associated with downregulated genes, and vice-versa, by HOMER.

RNA-seq analyses revealed genes that are differentially expressed in *FOXA-TKD* foregut cells (1792 up- and 767 down-regulated) (**Fig. 1B**). Down-regulated genes were enriched for foregut-related Gene Ontology (GO) terms, including “Morphogenesis of an epithelium”. Up-regulated/de-repressed genes were enriched for alternative mesodermal and ectodermal lineage GO terms (“Regulation of neuron differentiation”, “Angiogenesis”, “Muscle organ development”, “Epithelial to mesenchymal transition”) (**Fig. 1B**). Genes normally expressed at later stages of endoderm differentiation for liver and pancreas specification (e.g., *TBX3, PROX1*, and *SOX9*) were also de-repressed in the *FOXA-TKD* foregut cells. These genes were characterized by the GO term “Cell morphogenesis involved in differentiation”. RNA-seq analysis indicated that dox-induced *FOXA-TKD* in neuroectoderm cells had a negligible impact on global gene expression, where *FOXA* genes are not normally expressed (**Fig. S1D**). Thus, FOXA TFs are required for preventing precocious and alternative-lineage gene expression in the foregut endoderm, in addition to foregut gene activation.

To determine potential direct target genes of the FOXA TFs, we performed FOXA1 and FOXA2 CUT&RUN sequencing in control foregut cells. We identified 6115 combined FOXA1 and FOXA2 chromatin binding peaks (FOXA peaks) including those proximal to the known target genes, HNF1B and HNF4A (**Fig. S2A**). *De novo* motif analysis of FOXA peaks confirmed top enrichment of the FOXA binding motif, followed by known FOXA co-binding TF motifs (e.g., SOX, GATA, HNF1) (**Fig. 1C**). To associate FOXA peaks with potential direct target genes, we used the Genomic Regions Enrichment of Annotations Tool (GREAT) and identified 1167 FOXA peaks associated with 576 *FOXA-TKD*-mediated up-regulated genes (hypergeometric *P*-value = 1.6e-35) and 1011 FOXA peaks associated with 411 *FOXA-TKD*-mediated down-regulated genes (hypergeometric *P*-value = 2.47e-96) (*21*) (**Fig. 1D**).

FOXA’s role in gene activation is consistent with their canonical pioneering activity, but the mechanisms by which they directly repress gene expression during endoderm differentiation are unclear. To gain insight into these repressive mechanisms, we performed *de novo* motif analysis of FOXA peaks for the differentially expressed genes in *FOXA*-TKD. PR domain zinc finger 1 (PRDM1) TF-binding motifs were most significantly enriched at FOXA peaks associated with *FOXA*-TKD-mediated gene up-regulation (using FOXA peaks with *FOXA*-TKD-mediated gene down-regulation as background) (**Fig. 1E**). PRDM1 (BLIMP1) is a tumor suppressor and a repressor in primordial germ cell (*22*–*24*) and immune cell differentiation (*25, 26*). To adress the role of PRDM1 in endoermal differentiation, we generated a dox-inducible *PRDM1*-CRISPRi/KD foregut model and observed compromised foregut gene expression, with 249 up-regulated and 1397 down-regulated genes (**Fig. S2B**). 85% of the up-regulated genes were also up-regulated in *FOXA*-TKD foregut cells (212/249, hypergeometric *P*-value <1e-100), while 30% of the down-regulated genes were down-regulated in the *FOXA-TKD* foregut (420/1397, hypergeometric *P*-value <1e-100) (**Fig. 2A**).

**Fig. 2.**
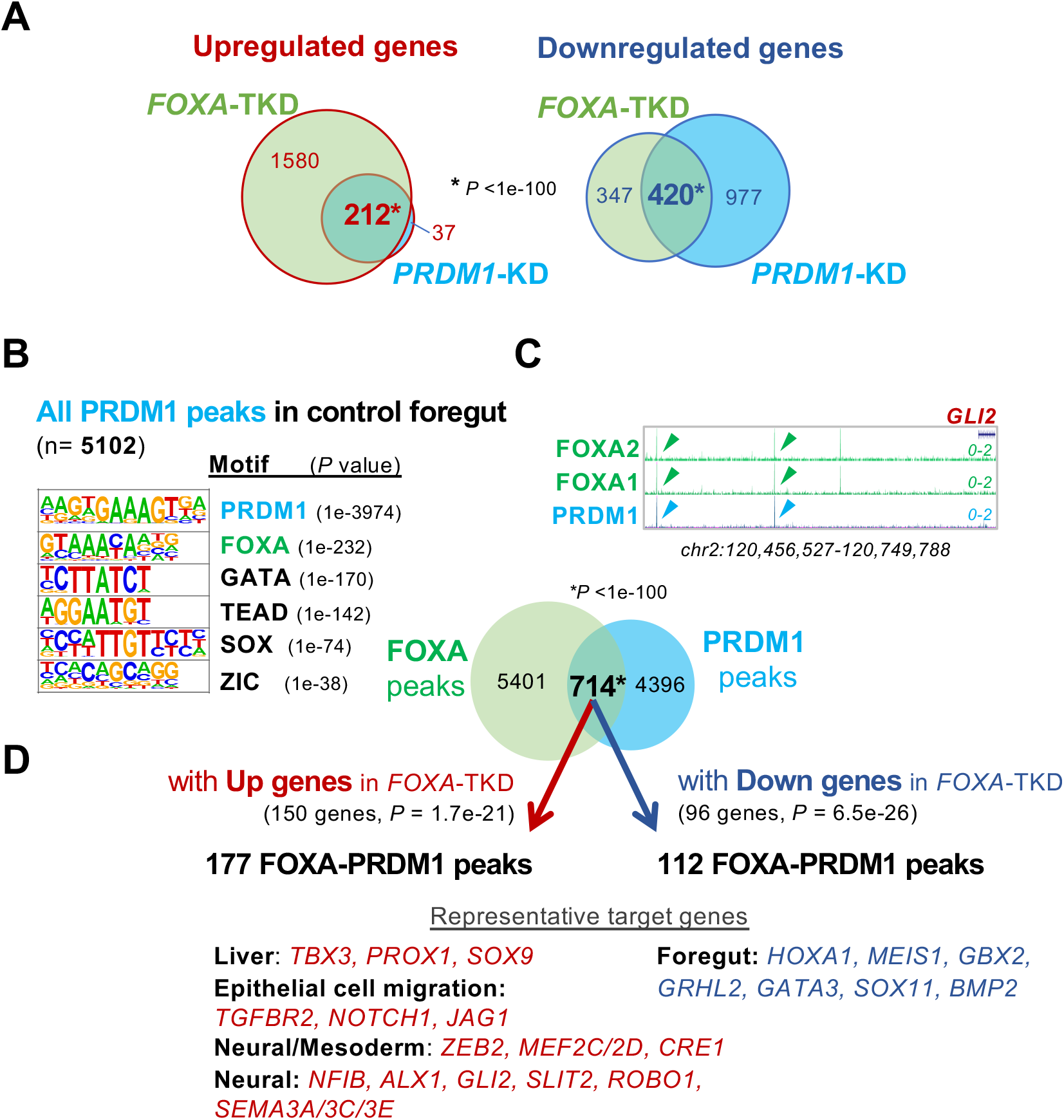
FOXA-PRDM1-bound enhancers prevent alternative-lineages and precocious gene expression while priming foregut gene activation. **(A)** Intersections of differentially expressed genes in *FOXA*-TKD and *PRDM1*-KD foregut cells. *P* value of the overlap by hypergeometric test. **(B)** Top motifs enriched at PRDM1 ChIP-seq peaks in control foregut cells by HOMER (n=2 replicates from 2 independent dox-untreated *FOXA*-CRISPRi clones). **(C)** FOXA1 and FOXA2 CUT&RUN and PRDM1 ChIP-seq tracks in control foregut cells at *GLI2* loci in RPM scale. Intersection of FOXA1/A2 CUT&RUN and PRDM1 ChIP-seq peaks in control foregut cells. *P* value of the overlap by hypergeometric test. **(D)** Intersections of FOXA-PRDM1 peaks associated with differentially expressed genes in *FOXA*-TKD foregut by GREAT. *P* value of the intersection by hypergeometric test. Representative target genes are shown.

Given the *FOXA genes* were down-regulated in *PRDM1*-KD foregut, the highly correlated gene expression changes in *FOXA*-TKD and *PRDM1*-KD could be a consequence of lower FOXA and PRDM1, with PRDM1 binding sites distributed independently of FOXA binding sites. To test this idea, PRDM1 ChIP-seq was performed in control foregut cells. *De novo* motif analysis of all 5102 PRDM1 binding peaks confirmed enrichment of the PRDM1 binding motif, followed by the FOXA binding motif (**Fig. 2B**). We found that PRDM1 peaks are significantly colocalized with FOXA peaks at over 700 genomic loci in foregut cells (**Fig. 2C**, hypergeometric *P*-value <1e-100), indicating that the PRDM1 binding sites are not distributed independently of FOXA binding sites and that the TFs likely function together in the same regulatory pathway.

To investigate this potential common regulatory function, we analyzed the binding sites and target genes in more detail. More than 95% of sites bound by both FOXA and PRDM1 are located outside of promoter regions (−1,000 bp to +100 bp of TSS) (**Fig. S2C**), suggesting that most sites are distal regulatory elements, i.e., enhancers. Among the 714 sites bound by both FOXA and PRDM1, 177 sites are associated with 150 de-repressed/up-regulated genes in *FOXA*-TKD foregut (**Fig. 2D**). The 150 de-repressed target genes include alternative-lineage regulators (neural and mesoderm genes), key liver TFs (e.g., *TBX3, PROX1*, and *SOX9*), and epithelial cell migration regulators (e.g., *TGFBR2, NOTCH1*, and *JAG1*) (**Fig. 2D**), which are normally up-regulated during later-stage liver differentiation.

Although PRDM1 motifs are preferentially enriched at FOXA peaks associated with *FOXA*-TKD-mediated up-regulated genes (**Fig. 1E**), we also found 112 FOXA-PRDM1–bound sites associated with 96 down-regulated genes in *FOXA-TKD* foregut (**Fig. 2D**). These differential transcriptional outcomes at FOXA-PRDM1 enhancers could be a consequence of divergent TF neighborhoods. The 112 down-regulated target genes include foregut-specific genes (e.g., *HOXA1, GBX2, GATA3*), which show basal expression levels in control cells from the hPSC to DE stages, followed by substantial up-regulation at the foregut stage (**Fig. S2D**). These results support a model in which FOXA-PRDM1–bound enhancers prevent precocious activation of late endoderm genes and alternative-lineage genes while priming the timely activation of foregut genes.

To examine epigenetic mechanisms by which the combinatorial binding of FOXA and PRDM1 might regulate both gene activation and repression, we analyzed the epigenetic features of all genomic sites bound by both FOXA and PRDM1 in foregut cells compared with “FOXA-only” and “PRDM1-only” bound sites (**Fig. 3A-E**), and then characterized the subset of “FOXA-PRDM1” bound sites associated with *FOXA*-TKD-mediated differentially expressed genes (**Fig. 3F**). Since the *FOXA* genes were markedly up-regulated during differentiation from definitive endoderm (DE) to foregut, we hypothesized that FOXA-PRDM1-dependent epigenetic features are established at the foregut stage. A time course of chromatin binding confirmed that FOXA2 and PRDM1 binding was substantially increased from the DE to foregut stage (**Fig. 3A**). ChIP-seq or CUT&RUN for two Polycomb repressive histone modifications (H2AK119ub1 and H3K27me3) and the active histone modification H3K4me1, revealed that H2AK119ub1 was substantially increased from the DE to foregut stage at FOXA-PRDM1–bound sites, but less so at FOXA-or PRDM1-only sites (**Fig. 3B**,). Similarly, the H3K27me3 Polycomb repressive mark was moderately and significantly increased in foregut at FOXA-PRDM1–bound sites (**Fig. 3B**). The H3K4me1 active mark was substantially increased from the DE to foregut stage at both FOXA-only and FOXA-PRDM1–bound sites (**Fig. 3B**), consistent with previous reports that FOXA interacts with a H3K4me1 methyltransferase (*27*). Altogether, our results revealed that the bivalent epigenetic states of Polycomb repressive marks and the H3K4me1 active mark are established at the foregut stage, correlated with the combinatorial binding of FOXA and PRDM1.

**Fig. 3.**
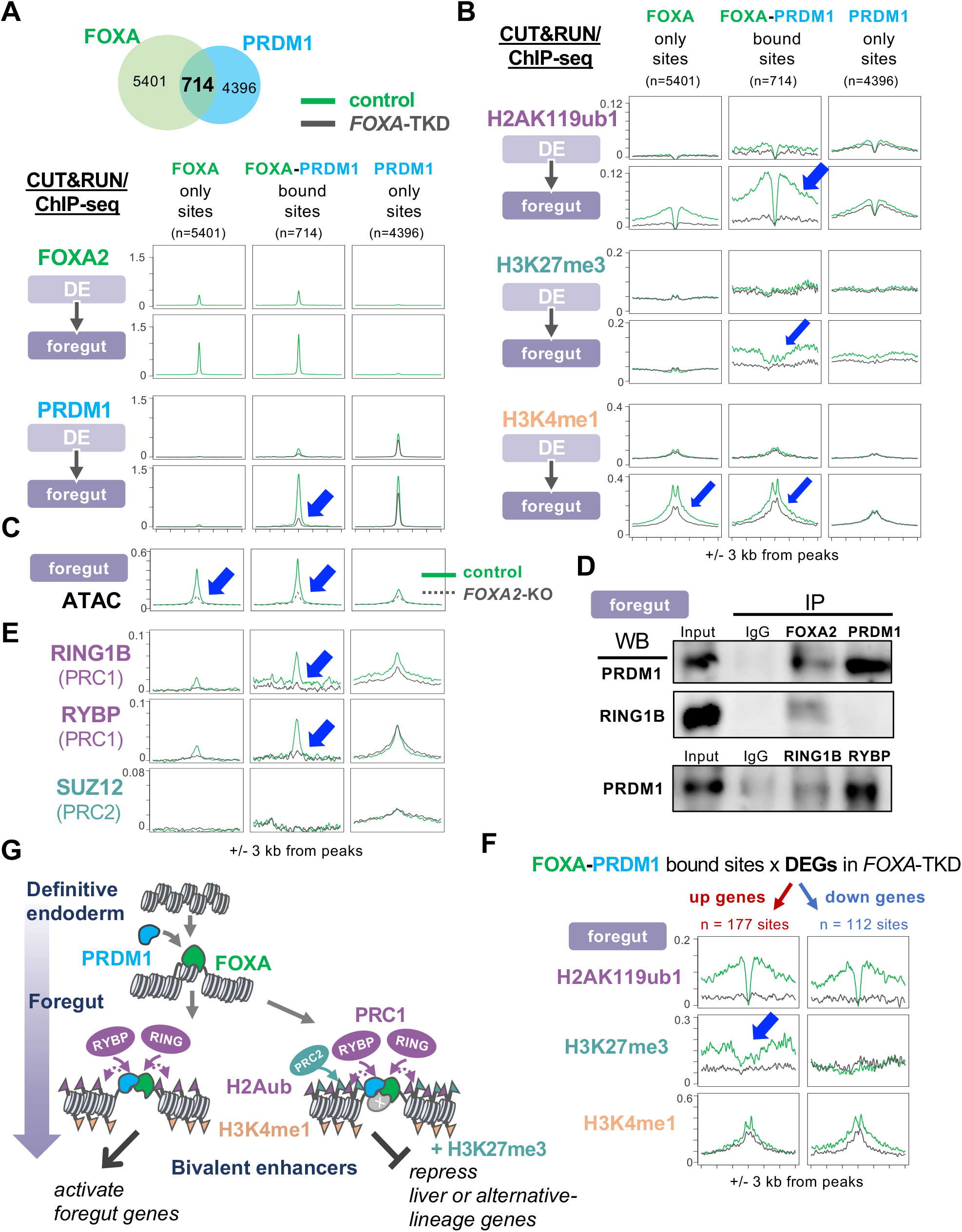
The combinatorial binding of FOXA and PRDM1 is required for establishing bivalent enhancers. **(A** and **B)** Averaged profiles of FOXA2 CUT&RUN, PRDM1 ChIP-seq, H2Aub ChIP-seq, H3K27me3 CUT&RUN, and H3K4me1 CUT&RUN in control (green lines) and *FOXA*-TKD (gray lines) definitive endoderm (DE) and foregut at FOXA-only bound sites, FOXA-PRDM1 bound sites, and PRDM1-only bound sites. **(C)** Averaged profiles of ATAC-seq in control (green lines) and *FOXA2* knockout (dotted gray lines) foregut cells. **(D)** co-immunoprecipitation (IP) western blot (WB) analysis between FOXA2, PRDM1 and PRC1 components (RING1B and RYBP). **(E)** Averaged profiles of RING1B, RYBP, and SUZ12 ChIP-seq in control (green lines) and *FOXA*-TKD (gray lines) foregut. **(F)** Averaged profiles of H2Aub, H3K27me3, and H3K4me1 in control (green lines) and *FOXA*-TKD (gray lines) foregut at FOXA-PRDM1 bound sites associated with differentially regulated genes in *FOXA*-TKD. **(G)** A model explaining the FOXA and PRDM1-mediated establishment of bivalent enhancers for bidirectional gene regulation. All CUT&RUN and ChIP-seq experiments, n=2 replicates from 2 independent *FOXA*-CRISPRi clones.

To determine whether FOXA TFs are involved in recruitment of PRDM1 and establishment of bivalent epigenetic states, we repeated ChIP-seq or CUT&RUN for PRDM1 and the epigenetic marks in *FOXA*-TKD cells. *FOXA*-TKD foregut cells displayed a significant loss of PRDM1 binding from FOXA-PRDM1–bound sites, and to a lesser extent at PRDM-only bound sites (**Fig. 3A, S3A, S4A**), consistent with FOXA-dependent recruitment of PRDM1. The minor reduction of PRDM1 binding signals at PRDM1-only sites in *FOXA-TKD* may be due to the 50% reduction in *PRDM1* expression observed in *FOXA-TKD* foregut, however, this is unlikely to account for the loss of PRDM1 chromatin binding at FOXA-PRDM1 bound sites.

Reduced PRDM1 binding at FOXA-PRDM1–bound sites could be due to loss of FOXA-mediated chromatin accessibility. As expected of pioneer factors, FOXA-bound sites (FOXA-only and FOXA-PRDM1) displayed decreased chromatin accessibility in *FOXA2* knockout (**Fig. 3C, S3C, S4C**) (ATAC from GSE114102) (*4*). Epigenetic analysis revealed that H2AK119ub1 and H3K27me3 were substantially decreased to background levels at FOXA-PRDM1–bound sites in *FOXA*-TKD foregut cells, whereas H3K4me1 was only modestly reduced (**Fig. 3B, S3B, S4B**). These results suggest that FOXA TF pioneering activity is required for the recruitment of PRDM1 to bivalent enhancers where they regulate Polycomb histone marks.

In addition to regulating chromatin accessibility, FOXA could directly recruit PRDM1 and/or Polycomb repressive complexes to lineage-specific enhancers. Co-immunoprecipitation (co-IP) of endogenous foregut proteins demonstrated physical interactions between FOXA2 and PRDM1, FOXA2 and the PCR1 subunit RING1B/RNF2, and PRDM1 and the PCR1 subunit RYBP (**Fig. 3D and S5A**). ChIP-seq revealed a significant loss of RING1B and RYBP binding from FOXA-PRDM1 bound sites in *FOXA*-TKD foregut cells relative to controls (**Fig. 3E, S3D, S4D**). By contrast, the PRC2 component SUZ12 showed low levels binding to FOXA-PRDM1–bound sites in control or *FOXA-TKD* foregut cells (**Fig. 3E**). Gene expression of PRC components was unaffected in *FOXA-TKD* foregut cells, consistent with a direct recruitment model. Collectively, these data suggest that FOXA promotes recruitment of PRDM1, which in turn coordinately recruits PRC1 (via interactions between FOXA and RING1B; PRDM1 and RYBP) and deposits Polycomb marks to establish *de novo* bivalent enhancers in the foregut endoderm.

To understand the differential transcriptional outcomes from FOXA-PRDM1–bound bivalent enhancers, we characterized the enhancers associated with differentially expressed genes in *FOXA*-TKD. The H3K27me3 repressive mark showed greater enrichment at FOXA-PRDM1 enhancers associated with *FOXA*-TKD-mediated gene up-regulation than at those with gene down-regulation, which was substantially decreased to background levels in *FOXA*-TKD (**Fig. 3F, S3E, S4E**). H2AK119ub1 and H3K4me1 marks were enriched to similar levels at both up-regulated and down-regulated FOXA-PRDM1 enhancers (**Fig. 3F, S3E**). These results suggest that H3K27me3 plays a repressive role, while H2AK119ub and H3K4me1 have dual or neutral roles in gene regulation.

PRCs and their marks have been mainly described as repressive, but evidence also suggests that PRC1/H2AK119ub1 support chromatin looping between enhancers and promoters to enable gene induction in response to activation signals (*28*–*31*). We propose two types of FOXA-PRDM1– mediated bivalent enhancers in normal foregut cells. The first enhancer type has Polycomb H2AK119ub1 and H3K27me marks, and H3K4me1 active mark, and mediates the repression of alternative-lineage and precocious gene expression. The second enhancer type has one Polycomb mark (H2AK119ub1) and the H3K4me1 mark and mediates the activation of foregut genes (**Fig. 3G**). We postulate that these differences in behaviors arise from additional TFs recruited to alternative-lineage enhancers in a FOXA-dependent manner to promote H3K27me3 deposition. To test this idea, we carried out d*e novo* motif analysis at FOXA-PRDM1 enhancers associated with *FOXA*-TKD-mediated gene up-regulation and identified binding motifs for the MSX TF (**Fig. S5B**), a repressor known to recruit PRC2 (*32*), which is up-regulated in foregut cells (**Fig. S1B**). We had previously identified the MSX binding motif at repressive FOXA binding sites in mouse endoderm (*33*), suggesting a conserved role for MSX in FOXA-mediated gene repression.

We asked whether our observations with FOXA and PRDM1 are specific to these factors or whether other pioneer TFs collaborate with PRDM TFs to regulate “appropriate” versus alternative-lineage genes. We examined the pluripotent pioneer TF, OCT4 (encoded by *POU5F1*), which has been reported to co-localize with 50% of the bivalent domains in hPSCs and interact with the PRC1 component RING1B/RNF2 in mouse PSCs (*16, 34*). It is unclear how OCT4 promotes active versus poised epigenetic states. Since PRDM14 has been reported to have overlapping binding pattern with Oct4 and form a complex with PRC2 in mouse pluripotent stem cells and with PRC1 in hPSCs (*35*–*40*), we tested whether OCT4 could coordinate with PRDM14 to regulate PRC recruitment and developmental genes.

We performed PRDM14 ChIP-seq in hPSCs and compared our data with publicly available OCT4 ChIP-seq peaks (*41*) (**Fig. 4A, S6A**). Consistent with our hypothesis, there was a significant overlap in OCT4 and PRDM14 chromatin binding in hPSCs (**Fig. 4A**, hypergeometric *P*-value <1e-100). *De novo* motif analysis of the 2067 PRDM14 binding peaks confirmed top enrichment of the PRDM14 binding motif, followed by the OCT4:SOX2 binding motif (**Fig. 4A**). 92% of genomic sites bound by OCT4 and PRDM14 (“OCT4-PRDM14” bound sites) are located outside of promoter regions (−1,000 bp to +100 bp of TSS) (**Fig. S6B**), suggesting that most of these sites are enhancers. OCT4-PRDM14 bound sites were enriched with Polycomb marks H2AK119ub1and H3K27me3, as well as the active mark H3K4me1, indicating a bivalent state (**Fig. 4B**, green plots). In contrast, OCT4-or PRDM14-only bound sites showed monovalent states, with either H3K4me1 or Polycomb marks, respectively (**Fig. 4B**). Polycomb marks were more enriched at OCT4-PRDM14–bound sites than at PRDM14-only sites, suggesting combinatorial recruitment of Polycomb repressive complexes by OCT4 and PRDM14.

**Fig. 4.**
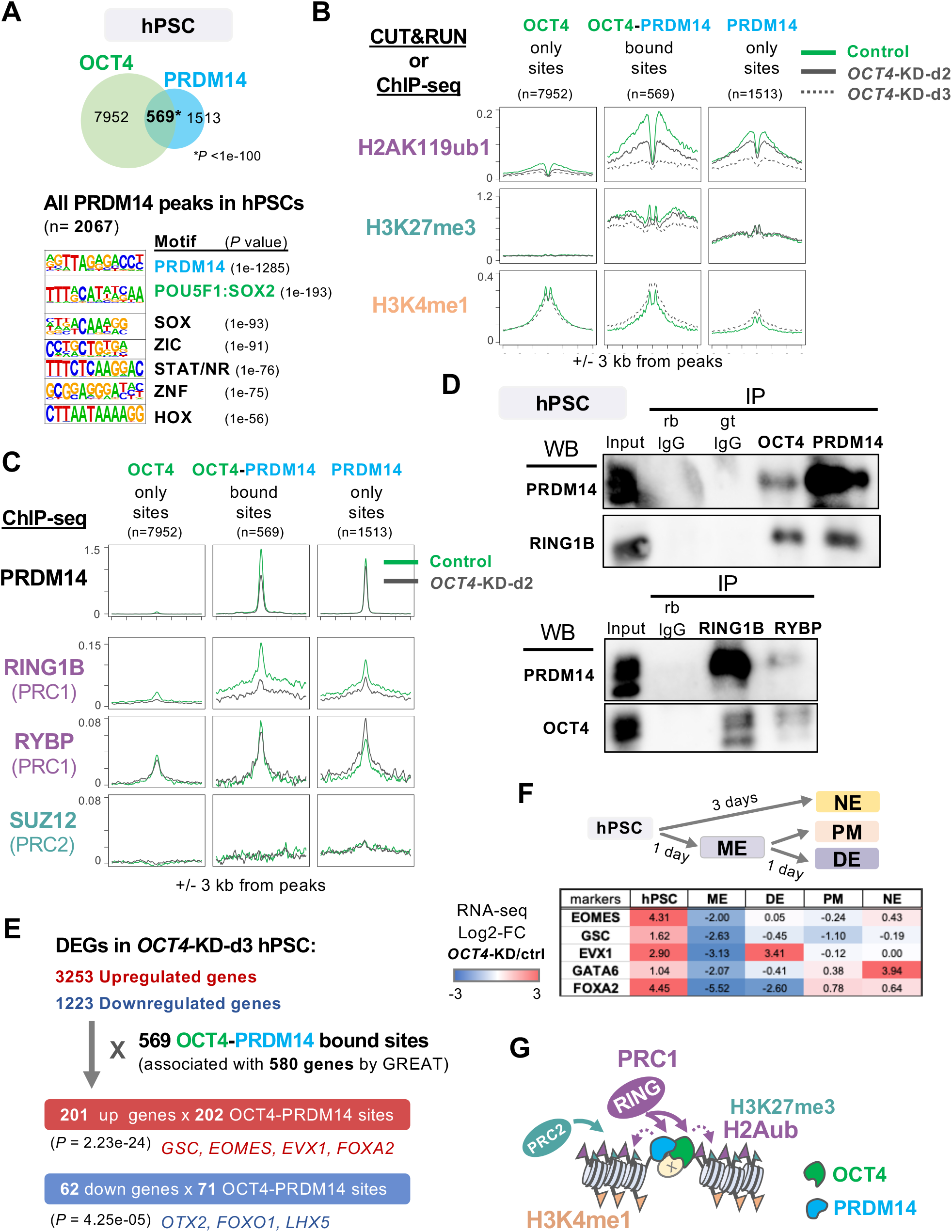
The combinatorial binding of OCT4 and PRDM14 recruits PRC1 to establish bivalent enhancers in hPSCs for poising early developmental genes. **(A)** Intersection of OCT4 and PRDM14 ChIP-seq peaks in control hPSCs. *P* value determined by hypergeometric test. Top motifs enriched at PRDM14 ChIP-seq peaks in control hPSCs by HOMER. **(B)** Averaged profiles of H2Aub ChIP-seq, H3K27me3 CUT&RUN, and H3K4me1 CUT&RUN in control (green lines), *OCT4*-KD-day2 (d2) (gray solid lines), and *OCT4*-KD-day3 (d3) (dotted gray lines) hPSCs at OCT4-only bound sites, OCT4-PRDM14 bound sites, and PRDM14-only bound sites. **(C)** Averaged profiles of PRDM14, RING1B, RYBP, and SUZ12 ChIP-seq in control (green lines) and *OCT4*-KD-day2 (d2) (gray lines) hPSCs. **(D)** co-immunoprecipitation (IP) western blot (WB) analysis between OCT4, PRDM14 and PRC1 components (RING1B and RYBP). **(E)** Intersections of OCT4-PRDM14 bound sites with differentially expressed genes (DEGs) in *OCT4*-KD hPSCs (n=3 replicates from 3 independent *OCT4*-CRISPRi clones; FDR < 0.05, FC > 1.5) by GREAT. Representative genes are shown. *P* value by hypergeometric test. **(F)** Log2 fold change (FC) of OCT4-PRDM14 target genes’ expression in *OCT4*-KD against control in hPSC (d2), mesendoderm (ME), definitive endoderm (DE), paraxial mesoderm (PM), and neuroectoderm (NE). **(G)** A model explaining the OCT4 and PRDM14-mediated establishment of bivalent enhancers. All CUT&RUN and ChIP-seq experiments, n=2 replicates from 2 independent *OCT4*-CRISPRi clones.

To assess whether OCT4 was required for enhancer bivalency, we generated dox-inducible *OCT4*-CRISPRi/KD hPSCs. No *OCT4* knock-down was detected at day-1 post dox, but by day-2 (*OCT4*-KD-d2), 80% mRNA and 50% protein were knocked down, and more than 95% mRNA and 85% protein were knocked down by day-3 post dox (*OCT4*-KD-d3) (**Fig. S6C**). H2AK119ub1 modifications at OCT4-PRDM14–bound sites were significantly and progressively reduced starting at *OCT4*-KD-d2, in parallel with the OCT4 reduction (**Fig. 4B, S7A, S8A**). By contrast, the H3K27me3 marks were only reduced at *OCT4*-KD-d3 (**Fig. 4B, S7A, S8A**), suggesting that H2AK119ub1 deposition is more sensitive to OCT4 levels than H3K27me3. The H3K4me1 mark was not affected even by *OCT4*-KD-d3 in hPSCs, indicating that, unlike FOXA, OCT4 is not required for H3K4me1 deposition (**Fig. 4B, S7A, S8A**).

To test whether OCT4 is required for recruiting PRDM14 and PRCs to OCT4-PRDM14– bound sites, we performed ChIP-seq of PRDM14 and PRC components in control and *OCT4*-KD-d2 hPSCs, where the expression of PRDM14 and PRC components were unaffected. We observed a significant loss of PRDM14 and RING1B/RNF2 (PRC1) ChIP-seq signals at OCT4-PRDM14– bound sites, and to a lesser extent at PRDM14-only bound sites (**Fig. 4C, S7B, S8B**). By contrast, RYBP (PRC1) and SUZ12 (PRC2) ChIP-seq signals were unaffected (**Fig. 4C, S7B, S8B**), suggesting their recruitment is OCT4-independent. Co-IP experiments of hPSC endogenous proteins supported physical interactions between OCT4, PRDM14, and RING1B (**Fig. 4D**). Collectively, our data suggest that OCT4 and PRDM14 cooperate to recruit the PRC1 component RING1B for H2AK119ub1 deposition at bivalent enhancers in hPSCs.

To determine the target genes of OCT4-PRDM14–bound bivalent enhancers, OCT4-PRDM14–bound sites were associated with differentially expressed genes in *OCT4*-KD hPSCs using GREAT (**Fig. 4E**). Among 569 OCT4-PRDM14–bound sites, 202 were associated with 201 up-regulated genes (hypergeometric *P*-value = 2.23e-24), and 71 sites were associated with 62 down-regulated genes (hypergeometric *P*-value = 4.25e-05) (**Fig. 4E**). The up-regulated genes in *OCT4*-KD hPSCs included key mesendoderm (ME) genes (*GSC, EOMES, EVX1*) and *FOXA2*, and the down-regulated genes included key pluripotent and neuroectoderm (NE) genes (*OTX2, FOXO1, LHX5*). Expression changes of these target genes in *OCT4*-KD were cell-culture context dependent (**Fig. 4F**). Although ME genes were de-repressed in *OCT4*-KD hPSCs, they were not fully induced in response to ME differentiation signal in *OCT4*-KD ME (**Fig. 4F**). These results suggest that OCT4-PRDM14–bound bivalent enhancers prevent spontaneous developmental gene activation in hPSCs while priming timely activation in response to differentiation cues.

Understanding how lineage-specific genes become transcriptionally primed while restricting alternative lineage programs provides insight into developmental competence in cell differentiation, regeneration, and tumorigenesis. The current dogma is that pioneer TFs establish primed and active epigenetic states to initiate new transcriptional programs. We revealed a role of the pioneer TFs FOXA and OCT4 (and by proxy, other pioneer TFs) in preventing inappropriate and/or precocious gene expression. This function is carried out by combinatorial binding of the pioneer and PRDM TFs followed by locus- and lineage-specific recruitment of PRC to establish bivalent enhancers. Our results highlight the distinctive mechanisms underlying PRC recruitment to enhancers versus to promoters, the latter of which has been extensively studied and mainly explained by more generic targeting mechanisms (*9*) We postulate that FOXA TFs take advantage of available corepressors, which contribute to lineage-specific gene repression depending on target genes and biological contexts. For example, the co-binding of Foxa2 and Rfx1 represses Cdx2 expression in adult mouse liver (*42*), and Foxa recruits Grg3/TLE3 to establish stably closed chromatin domains in hepatocytes (*43*). The role of pioneer TF-mediated gene repression likely extends to diverse biological contexts and could be applied to improve the efficiency and integrity of cellular reprogramming.

## Acknowledgments

We thank D. Haslam, B. Gebelein, and A. Barski for sharing equipment and materials; Y-C. Hu, J. Tchieu, and K, Loh for technical advice; B. Gebelein, V. Hwa, A. Zorn, R. Kopan, K. Zaret, G. Mirizio, G. Riddihoug for helpful comments and editing on the paper; the Digestive Disease Research Core Center in CCHMC (Pluripotent Stem Cell Facility, Genome Edition Core, DNA sequencing Core, Research Flow Cytometry Core, Confocal Microscopy Core, Viral Vector Production Core).

## Funding

Cincinnati Children’s Research Foundation, Trustee Award to MI, HL

Cincinnati Children’s Research Foundation, Center for Pediatric Genomics Pilot Award to HL

National Institutes of Health, P30 DK078392 to MI

National Institutes of Health, 1R01GM143161 to MI

Japan Society for the Promotion of Science Foundation, Postdoctoral Fellowship to SM Uehara Memorial Foundation, Postdoctoral Fellowship to SM

## Author contributions

Conceptualization and Supervision: MI

Investigation: SM, MG, MI, MB

Software and statistical analysis: HL

Bioinformatic analysis: HL, MI

Resources: JS, CM, WZ

Writing – Original Draft: MI

Writing – Review & Editing: All authors

Funding acquisition: MI, HL

## Supplementary Materials for

## Materials and Methods

### Human induced pluripotent stem cell (hPSC) culture

Human induced pluripotent stem cells (72.3, male, RRID: CVCL_A1BW) were obtained from the CCHMC Pluripotent Stem Cell Facility (PSCF) (*44*) and cultured in feeder-free condition with mTeSR1 (85850, StemCell Technologies) and on a hESC-qualified Matrigel (354277, Corning) or Cultrex Stem Cell Qualified Reduced Growth Factor Basement Membrane Extract (3434-010-02, Biotechne) coated plate. Spontaneously differentiated cells were manually removed before passage and differentiation. For passaging, hPSC colonies were dissociated with Gentle Cell Dissociation Reagent (07174, StemCell Technologies) into small cell clusters and seeded on Matrigel or Culturex coated culture plates and cultured in mTeSR1. 72.3 iPSCs were distributed by the PSCF from a quality controlled, cryopreserved bank prepared at ∼p30. Cells were documented to be mycoplasma-free, harbor a normal G-banded karyotype, and identity was confirmed by STR profiling.

### Generation of dox-inducible CRISPRi hPSC line

Dox-inducible CRISPRi host hPSCs were established essentially as previously described (*19*). Plasmid pAAVS1-NDi-CRISPRi (Gen 1) was a kind gift of Dr. Bruce Conklin (73497, Addgene plasmid). The gRNA sequence for targeting the AAVS1 locus (sgRNA-T2: GGGGCCACTAGGGACAGGAT) (*45*) was subcloned into the pX459M2-HF vector, a modified version of the pX459 V2.0 vector (62988, Addgene plasmid) carrying an optimized gRNA scaffold, to generate plasmid pX459M2-HF-AAVS1(*46, 47*). Briefly, 72.3 cells were reverse transfected (TRANS-LT1, Mirus) with 2ug each of pX459M2-HF-AAVS1 and pAAVS1-NDi-CRISPRi (Gen 1). Clones that formed during 8 days of exposure to 100μg/mL G418 were manually excised, expanded, and genotyped. A clone containing bi-allelic knock-in of a cassette for constitutive expression of the third-generation modified rtTA protein (Tet-On 3G) and doxycycline inducible KRAB-dCas9-P2A-mCherry into the AAVS1 locus was identified by Sanger sequencing and used for subsequent experiments. Dox-inducible CRISPRi host hPSCs were distributed by the PSCF from a quality controlled, cryopreserved bank prepared at ∼p50, transduced gRNA (see below), and used in experiments up to p72 (*FOXA*-CRISPRi), p64 (*OCT4*-CRISPRi), and p79 (*PRDM1*-CRISRPi). Cells were documented to be mycoplasma-free, harbor a normal G-banded karyotype, and identity was confirmed by STR profiling.

### gRNA Design and Cloning into the gRNA-Expression Vector

gRNAs targeting *FOXA1/A2/A3* loci (for *FOXA*-CRISPRi/TKD), *PRDM1* locus (for *PRDM1*-CRISPRi/KD), or *OCT4* locus (for *OCT4*-CRISPRi/KD) were designed by a comprehensive algorithm to predict high efficiency for CRISPRi (*48*) and listed below. Two to four gRNAs per each CRISPRi target drived from independent RNA polymerase III promoters were cloned into a single lentivirus expression vector by a two-step Golden Gate cloning method, as previously described (*49*). Briefly, at step 1, annealed and phosphorylated oligos for each desired genomic target were cloned into pmU6-gRNA (53187, Addgene), phU6-gRNA (53188, Addgene), phH1-gRNA (53186, Addgene) or ph7SK-gRNA (53189, Addgene) plasmid by Golden gate assembly method. At step 2, plasmids from the step 1 were cloned into pLV GG hUbC-EGFP (84034, Addgene; dsRED was replaced with EGFP from 1168, Addgene) by Golden gate assembly method.

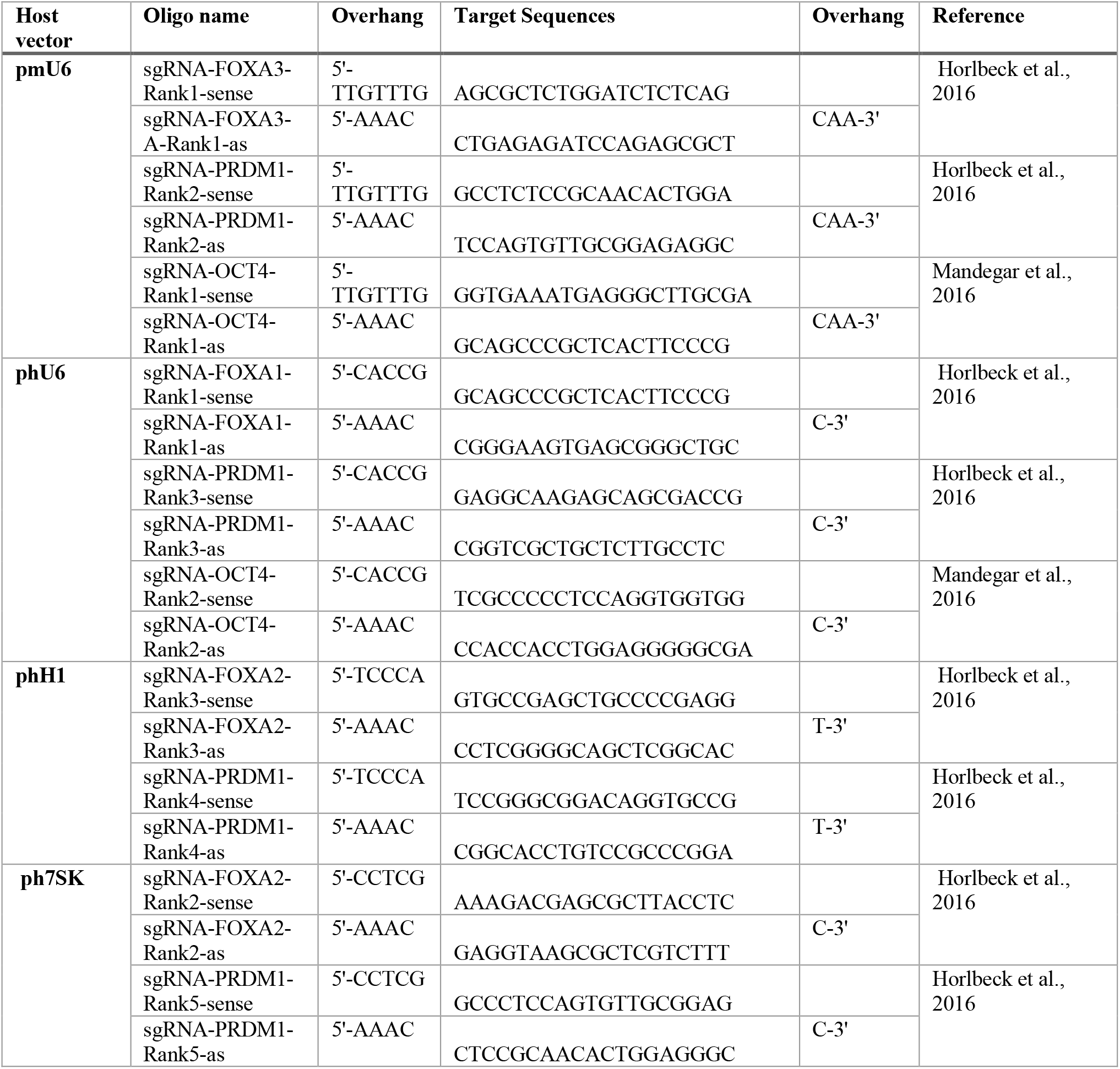

### Viral production, transduction, and establish FOXA-CRISPRi, PRDM1-CRISPRi, and OCT4-CRISPRi monoclonal lines

The lentivirus was produced by the CCHMC viral vector core or in house. Briefly, gRNA-expression lentivirus vector, packaging plasmid (psPAX2) and envelope plasmid (pMD2.G) were transfected to HEK293T cell by the polycation polyethylenimine (PEI) transfection method. The viral supernatant was passed through a 0.45 um filter and concentrated to 1:66 with Lenti-X concentrator (631231, Takara Bio) in house or to 1:255 by ultracentrifugation in the core.

Each Lentivirus was transduced to CRISPRi hPSCs by spin infection method (*50*). CRISPRi hPSCs were dissociated to single cells via Accutase, the dissociated cells were resuspended in the transduction medium (1 uL to 5 uL of lentivirus, 4 to 8 ug/ml Polybrane[STR-1003-G, Sigma Aldrich], and 10 uM Y27632 [72304, StemCell Technologies] in mTeSR1), and centrifuged at 3200 xg for 30 to 90 min. The cell pellet was resuspended in mTeSR1 supplemented with 10 uM Y27632 and plated on the Matrigel coated plate. Next day, culture media was changed to mTeSR1, and daily maintenance was performed until sub-confluence. EGFP-positive lentivirus transduced cells were sorted by Moflo XDP (Beckmann Coulter) or BD FACSAria II (BD Biosciences) and cloned from single cells by limiting dilution method. Clonal cells were cultured with mTeSR1 supplemented with 1x CloneR (StemCell Technologies) for 2 days. At day 3, 25% volume of 1x CloneR-supplemented mTeSR1 was added. At day 4 and after, culture media was changed to mTeSR1, and daily maintenance was performed. After expansion of monoclonal lines, a minimum of 6 clones were tested for CRISPRi knockdown efficiency by RT-qPCR. To induce CRISPRi knockdown of target genes, cells were cultured in the presence of 500 nM to 2 uM doxycycline (D9891-5G, Sigma-Aldrich).

### hPSC to Endoderm, Mesoderm, and Neuroectoderm differentiation

For hPSC differentiation protocols, hPSCs were dissociated as single cells with Accutase (07922, StemCell Technologies; A6964-500ML, Sigma-Aldrich). Dissociated cells were plated on a Matrigel or Culturex coated culture plate and cultured in 10 uM Y27632 (72304, StemCell Technologies) containing mTeSR1 one day before differentiation. Chemically defined basal media 2 (CDM2) and CDM3 were composed as previously reported (*51*), filtered through 0.22 um unit, and stored at -80°C until use. Once thawed, CDM2 was used within 2 weeks, and CDM3 was used within 3 weeks. Each differentiation protocol is shown below.

### Endoderm differentiation

Endodermal induction was performed essentially as previously described (*20, 51*). For mesendoderm (ME) induction, hPSCs were cultured in CDM2 supplemented with 100 ng/ml ActivinA (800-0, Shenandoah; or GFH6, Cell Guidance Systems), 2 uM CHIR99021 (SML1046-25MG, Sigma-Aldrich), and 50 nM PI103 (2930-1, Tocris Bioscience) for 24 hours. For definitive endoderm (DE) induction, ME cells were briefly washed with DMEM/F12 and cultured with CDM2 supplemented with 100 ng/ml ActivinA and 250 nM LDN193189 (SML0559-5MG, Sigma-Aldrich) for 24 hours. For foregut endoderm (FG) induction, DE cells were briefly washed with DMEM/F12 and cultured with CDM3 supplemented with 1 uM A83-01 (SML0788-5MG, Sigma-Aldrich), 2 uM ATRA (R2625-100MG, Sigma-Aldrich), 10 ng/ml bFGF (PHG0261, Thermo Fisher Scientific), and 30 ng/ml BMP4 (314-BP-050, R&D systems) for 24 hours. For Liver bud (LB) induction, FG cells were cultured in CDM3, supplemented with 10 ng/ml Activin A, 30 ng/ml BMP4, 1 μM Forskolin (F3917-10MG, Sigma-Aldrich) for 72 hours. Differentiation media was changed every 24 hours.

### Paraxial mesoderm differentiation

Paraxial mesodermal induction was performed with some modification of the published protocol (*52*). For mesoendoderm (ME) differentiation, hPSCs were cultured in CDM2 supplemented with 100 ng/ml ActivinA, 2 uM CHIR9902, and 50 nM PI-103 for 24 hours. For paraxial mesoderm induction, ME cells were briefly washed with DMEM/F12 and cultured in CDM2 supplemented with 1 uM A83-01, 2 uM CHIR99021, and 250 nM LDN193189 for 24 hours.

### Neuroectoderm differentiation

hPSCs were cultured in CDM2 supplemented with 10% KOSR (10828028, ThermoFisher Scientific), 250 nM LDN193189, 10 uM SB431542 (S4317-5MG, Sigma-Aldrich), and 5 uM XAV931 (X3004-5MG, Sigma-Aldrich) for 3 days. Differentiation media was changed with every day.

### RT-qPCR

500 ng of total RNA was reverse transcribed with iScript™ cDNA Synthesis Kit (1708890, BioRad). The synthesized cDNA was diluted in 1:5 ratio with water for RT-qPCR. SYBR Green method (A25742, ThermoFisher Scientific) was used for RT-qPCR reaction. *GAPDH* was used as an internal control. RT-qPCR reaction was run in QuantStudio™ 3 Real-Time PCR System (ThermoFisher Scientific). Relative gene expression was calculated by 2^-ΔΔCt^ method. Primer sequences were listed below.

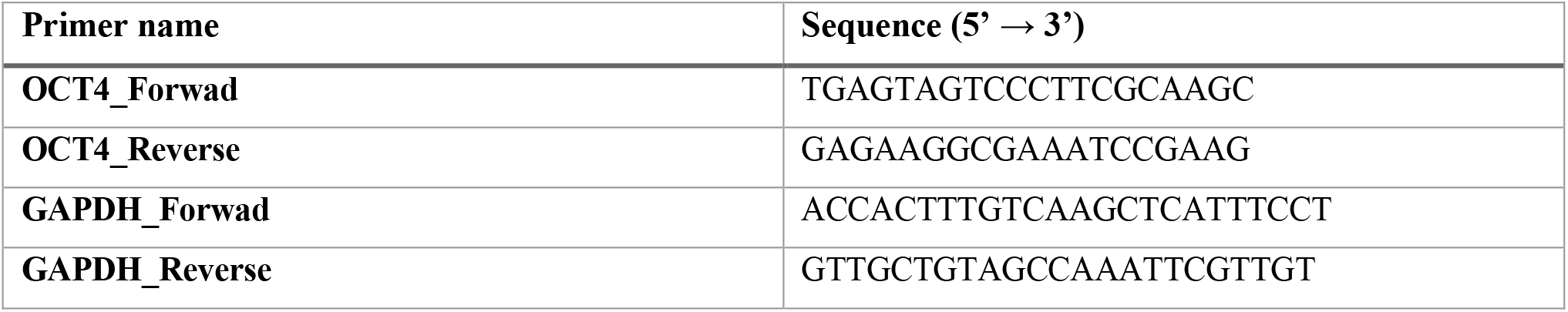

### Western Blot Assay

1 to 1.5 ug of nuclear protein in Laemmli Sample Buffer (J60015, Alfa Aesar) was denatured at 100°C for 5 min, run on 10 or 12.5% SDS-PAGE gel at 60 V for 30 min and then 100 V for 2 hours, and transferred to 0.2 or 0.45 um pore size PVDF membrane (LC2002, ThermoFishir Scientific and IPSN07852, Millipore) in wet tank transfer system with Mini Blot Module (ThermoFishir Scientific) at 20 V for 1 hour. The transferred membrane was briefly washed with TBST (0.1% Tween-20 in TBS) and blocked with Blocking Buffer (5% Blotting-Grade Blocker [1706404, BioRad] or 5% BSA [A9647-50G, Sigma-Aldrich] in TBST) for 1 hour at room temperature. The blocked membrane was washed with TBST at room temperature and incubated with primary antibody diluted with Blocking Buffer at 4°C overnight. Next day, the membrane was washed three to five with TBST for 5 min at room temperature and incubated with secondary antibody, diluted with 1% BSA in TBST for 1 hour at room temperature. The membrane was washed three to five with TBST for 5 min at room temperature. Chemiluminescence signal was developed with ECL Select Western Blotting Detection Reagent (RPN2235, GE Healthcare) or Clarity™ Western ECL Substrate (170-5061, BioRad) for 5 min. Chemiluminescence image was acquired by ChemiDoc™ MP imaging system (Bio-Rad). Antibodies used in Western Blot Assay were listed in below.

| Antigen | Supplier | Cat. No. | Lot No. | Host | Dilution factor |
| --- | --- | --- | --- | --- | --- |
| FOXA1 | Abcam | ab55178 | GR3283617-4 | Mouse | 1:1000 |
| FOXA2 | Abnova | H00003170-M01 | IA081-7E6 | Mouse | 1:1000 |
| PRDM1 | CST | 9115S | 6 | Rabbit | 1:500 |
| OCT4 | R&D systems | AF1759 | JTW0419081 | Goat | 1:1000 |
| PRDM14 | Abcam | ab187881 | GR3297404-8 | Rabbit | 1:1000 |
| RING1B | BETHYL | A302-869A | 1 | Rabbit | 1:1000 |
| **RYBP** | Millipore | AB3637 | 3123446 | Rabbit | 1:1000 |
| **VINCULIN** | Sigma-Aldrich | V-9131 | 118M4777V | Mouse | 1:5000 |
| **Anti-Mouse IgG_HRP** | GE Healthcare | NA931 | 11959045 | Sheep | 1:5000 |
| **Anti-Rabbit IgG_HRP** | GE Healthcare | NA934 | 11985050 | Donkey | 1:5000 |
| **TrueBlot Anti-Goat IgG_HRP** | ROCKLAND | 18-8814-31 | 46356 | Mouse | 1:1000 |
| **TrueBlot Anti-Rabbit IgG_HRP** | ROCKLAND | 18-8816-33 | 43708 | Mouse | 1:1000 |

### Co-Immunoprecipitation (co-IP)

We modified published co-IP protocols (*53, 54*). Cultured cells were briefly washed with DMEM/F12 and crosslinked with 1.5 mM DSP (22585, ThermoFisher Scientific) in PBS at room temperature for 30 min with gentle shaking. The crosslinking solution was aspirated, and crosslinking reaction was quenched with 30 mM Tris-HCl (pH 7.4) in PBS at for 20 min with gentle shaking. Crosslinked cells were briefly washed twice with ice-cold 1x PBS and scraped in ice-cold 0.01% PVA in 1x PBS. The cell suspension was transferred to a conical tube and centrifuged at 350 xg for 3-5 min at 4°C. Supernatant was removed, and cell pellet was re-suspended with ice-cold 0.01% PVA in 1x PBS and centrifuged at 800 xg for 3-5 min at 4°C. Supernatant was removed, and cell pellet was snap frozen by dry ice. Cell pellet was stored at - 80°C until going on the next step.

For nuclear protein extraction, cell pellet was suspended with Cell Lysis Buffer (5 mM HEPES-NaOH [pH. 7.9], 10 mM KCl, 1 mM DTT, 0.5% NP-40, 1x protease inhibitor) and incubated on ice for 10 min. Then, cell suspension was centrifuged at 1700 xg for 10 min at 4°C, and supernatant was removed. Cell pellet was re-suspended with Nuclear Lysis Buffer (100 mM NaCl, 25 mM HEPES-NaOH [pH. 7.9], 1 mM MgCl_2_, 0.2 mM EDTA, 0.5% NP-40, 600U/mL Benzonase [70664, Millipore], 1x protease inhibitor) and incubated for 4 hours at 4°C with rotation. NaCl concentration of nuclear lysate was adjusted to 200 mM and incubated for 30 min at 4°C with rotation. Nuclear lysate was centrifuged at maximum speed (16000 xg) for 30 min at 4°C. Take appropriate volume of precleared lysate as an input. Incubate lysates with appropriate amount of antibody (listed below) overnight at 4°C with rotation. For each immunoprecipitation (IP) reaction, 50 uL Dynabeads Protein-G beads (10004D, Thermo Fisher Scientific) was used. Dynabeads were washed twice with PBST (0.1% Tween-20 in PBS). Washed Dynabeads were resuspended with antibody-cell lysate and incubated for 2 hours at 4°C with rotation. Dynabeads were washed 5 times with IP-Wash Buffer (100 mM NaCl, 25 mM HEPES-NaOH [pH. 7.9], 1 mM MgCl_2_, 0.2 mM EDTA, 0.5% NP-40, 1x protease inhibitor) at room temperature for 2 min with rotation. Dynabeads-bound antibody-proteins (IP samples) were eluted with 1x Laemmli Sample Buffer (1610737EDU, Bio-Rad) for 20 min at 65°C while shaking at 1000 rpm in ThermoMixer F1.5 (Eppendorf). For denaturing IP and Input samples, add 1/20 volume of 2-mercaptoethanol and boil for 5 min at 98°C. Denatured IP and Input samples were stored at -20°C and used for Western Blot assay.

| **Antigen** | **Supplier** | **Cat. No.** | **Lot No.** | **Host** | **Antibody amount** |
| --- | --- | --- | --- | --- | --- |
| **FOXA2** | Millipore | 07-633 | 2971828 | Rabbit | 5.0 ug |
| **PRDM1** | CST | 9115S | 6 | Rabbit | 0.1 ug |
| **OCT4** | R&D systems | AF1759 | JTW0419081 | Goat | 1.0 ug |
| **PRDM14** | Abcam | ab187881 | GR3297404-8 | Rabbit | 5.0 ug |
| **RING1B** | BETHYL | A302-869A | 1 | Rabbit | 4.0 ug |
| **RYBP** | Millipore | AB3637 | 3123446 | Rabbit | 5.0 ug |

### RNA Sequencing

1 million cultured cells were dissociated to single cells with Accutase and were mixed with 100,000 of *Drosophila* Schneider 2 (S2) cells for a spike-in control. Cells were pelleted and lysed with Aurum Total RNA Lysis Solution (BioRad # 7326802). Total RNA was extracted by Aurum™ Total RNA Mini Kit (BioRad #7326820) and sent to Novogene for library preparation and paired-end-150bp sequencing on an Illumina NovaSeq 6000.

### Chromatin Immunoprecipitation (ChIP)

Cells were crosslinked with 1% formaldehyde in 1x PBS for 10 min at room temperature. The crosslinking reaction was stopped by adding 0.125 M glycine for 5 min at room temperature. Crosslinked cells were centrifuged at 600 xg for 5 min at 4°C, and supernatant was removed. The cell pellet was washed with ice-cold 1x PBS and centrifuge at 600 xg for 5 min at 4°C. Repeat the wash step (total twice). The cell pellet was frozen by dry ice and stored at -80°C.

Crosslinked human cells were resuspended with Lysis Buffer1 (10 mM Tris-HCl [pH. 8.0], 10 mM NaCl, 0.5% NP-40, and 1x Complete Protease Inhibitor EDTA free) and incubated for 10 min on ice. 25% crosslinked *Drosophila* S2 cells were reconstituted with Lysis Buffer1as a spike-in control and was mixed with the human cells. Cells were centrifuged at 665 xg for 5 min at 4°C, and supernatant was removed. Cell pellet was resuspended with Lysis Buffer2 (50 mM Tris-HCl [pH. 8.0], 10 mM EDTA, 0.32% SDS, 1x Complete Protease Inhibitor EDTA free) and incubated for 10 min on ice. Cell lysate was diluted with IP Dilution Buffer (20 mM Tris-HCl [pH. 8.0], 150 mM NaCl, 2 mM EDTA, 1% TritonX-100, 1x Complete Protease Inhibitor EDTA free) and transferred into a milliTUBE ATA Fiber (Covaris). Cell lysate was sonicated by Covaris S220 (Covaris) for 3.5 min. The insoluble debris was removed by centrifuge at 12000 xg for 5 min at 4°C. Sonicated chromatin (supernatant) was transferred to a new tube and stored at -80°C.

50 to 86 uL Dynabeads Protein-G beads (10004D, Thermo Fisher Scientific) were washed twice with PBS-T (0.02% Tween-20 in 1x PBS). Desired amounts of antibodies (listed below) were conjugated to washed beads for 2 to 6 hours at 4°C with rotation. Meanwhile, Frozen chromatin was thawed on ice. Desired amount of chromatin (listed below; determined by DNA amount) was diluted with IP dilution buffer in 4:1 ratio. Antibody conjugated Dynabeads were washed twice with IP Dilution Buffer (without Complete protease inhibitor EDTA free). Washed Dynabeads were resuspend with diluted chromatin and incubated for overnight at 4°C with rotation. Next day, ChIPed beads were washed four times with following washing buffers: FA Lysis Buffer (50 mM HEPES-KOH [pH. 7.5], 150 mM NaCl, 2 mM EDTA, 1% TritonX-100, 0.1% Sodium deoxycholate, 1x Complete protease inhibitor EDTA free); NaCl Buffer (50 mM HEPES-KOH [pH. 7.5], 500 mM NaCl, 2 mM EDTA, 1% TritonX-100, 0.1% Sodium deoxycholate); LiCl Buffer (100 mM Tris-HCl [pH. 8.0], 500 mM LiCl, 1% NP-40, 1% Sodium deoxycholate); and 10 mM Tris-HCl [pH. 8.0]. Chromatin was eluted with TES (50 mM Tris-HCl [pH. 8.0], 10 mM EDTA, 1% SDS) for 15 min at 65°C while shaking at 1000 rpm in ThermoMixer F1.5 (Eppendorf). The ChIP and Input chromatin were reverse crosslinked by adding NaCl (Final concentration of 200 mM) for 2 to 8 hours at 65°C. The reverse crosslinked chromatin samples were treated with 50 ng/ul RNaseA (EN0531, ThermoFisher Scientific) for 0.5 to 1 hour at 37°C. Samples were treated with 0.2 mg/ml protease K (03115828001, Roche) for 2 to 4 hours at 37°C. Finally, DNA was purified by phenol-chloroform extraction followed by ethanol precipitation. The DNA concentration was measured by Quantus fluorometer (Promega).

| Antigen | Supplier | Cat. No. | Lot No. | Host | Antibody amount [ug] | Chromatin (DNA) amount [ug] |
| --- | --- | --- | --- | --- | --- | --- |
| PRDM1 | CST | 9115S | 6 | Rabbit | 0.1 ug | 75 ug |
| PRDM14 | Abcam | ab187881 | GR3297404-8 | Rabbit | 11.5 ug | 37.5 ug |
| RING1B | BETHYL | A302-869A | 1 | Rabbit | 3.0 ug | 37.5 ug |
| RYBP | Millipore | AB3637 | 3123446 | Rabbit | 4.0 ug | 37.5 ug |
| SUZ12 | CST | 3737S | 8 | Rabbit | 0.42 ug | 37.5 ug |
| H2AK119Ub1 | CST | 8240S | 8 | Rabbit | 2.5 ug | 18.75 ug |

### CUT&RUN

20 uL of ConcanabalinA coated beads (BP531, Bangs Laboratories) were washed twice with Binding Buffer (20 mM HEPES-KOH [pH. 7.9], 10 mM KCl, 1 mM CaCl_2_ and 1mM MnCl_2_). Dissociated live cells were resuspended with Wash Buffer (20 mM HEPES-NaOH [pH. 7.5], 150 mM NaCl, 0.5 mM Spermidine and 1x Complete Protease Inhibitor EDTA free) and mixed with washed beads. The mixture was rotated for 10-15 min at room temperature. We then resuspend cell-bound beads with 100 ul (for histone mark CUT&RUN) and 50 ul (for TF CUT&RUN) Antibody Buffer (2 mM EDTA in Digitonin Buffer [0.02% Digitonin in Wash Buffer]). Desired amounts of antibodies (listed below) were added to cell-beads suspension and incubate overnight at 4°C with rotation. Next day, beads were washed twice with Digitonin Buffer. Washed beads were treated with 700 ng/ml pA-MNase in Digitonin Buffer for 1 hour at room temperature with rotation. Beads were washed twice with Digitonin Buffer. Washed beads were resuspended with 100 ul of Digitonin Buffer and place on the cool block (BP361036, Fisher Scientific) for >10 min to cool down to 0°C. We then performed pA-MNase digestion by adding 2 ul of ice-cold 0.1 M CaCl2 and incubating for 30 min on the cool block. To stop pA-MNase reaction, 100 ul of 2x Stop Buffer (340 mM NaCl, 20 mM EDTA, 4 mM EGTA, 0.02% Digitonin, 0.05 mg/ml RNaseA and 0.05 mg/ml Glycogen) was added. To release CUT&RUN fragment, samples were incubated at 37°C for 30 min with shaking at 300 rpm in ThermoMixer F1.5 (Eppendorf). The beads were removed by centrifuge at 16000 xg for 5 min at 4°C and placing on a magnetic stand. Supernatant was transferred and treated with 0.1% SDS and 0.25 mg/ml ProteinaseK for 1 hour at 37°C. Finally, DNA was purified by phenol-chloroform extraction followed by ethanol precipitation. The DNA concentration was measured by Quantus fluorometer (Promega).

| Antigen | Supplier | Cat. No. | Lot No. | Host | Antibody amount | Cell number |
| --- | --- | --- | --- | --- | --- | --- |
| FOXA1 | Abcam | ab55178 | GR3283617-4 | Mouse | 2.0 ug | 5.0x10<sup>5</sup> cells |
| FOXA2 | Millipore | 07-633 | 2971828 | Rabbit | 5.0 ug | 5.0x10<sup>5</sup> cells |
| H3K4me1 | Millipore | 07-436 | 3120573 | Rabbit | 2.0 ug | 2.5x10<sup>5</sup> cells |
| H3K27me3 | CST | 9733S | 14 | Rabbit | 0.102 ug | 2.5x10<sup>5</sup> cells |

### Preparation and sequencing of ChIP-seq and CUT&RUN-seq libraries

We prepared multiplexed libraries of ChIP, Input for ChIP, and CUT&RUN samples from two replicate (two independent CRISPRi clones) using NEBNext Ultra II DNA Library Prep Kit for Illumina (E7645, NEB). The manufacture’s protocol was employed with the following changes. After the adaptor ligation, we performed cleanup by AMPure XP Beads (A63880, Beckman Coulter) or MagBind Magnetic Beads (M1378-01, Omega) without size selection. After the RCR enrichment of adaptor-ligated library, we performed two rounds of size selection by the magnetic beads.

### RNA-seq data analysis

Sequencing reads were aligned to hg38 and dm6 combined genome using STAR aligner (*55*). Only uniquely and concordantly aligned read pairs were used for downstream analysis. Read count mapped to each gene was measured using featureCounts (*56*) with options ‘-O --fracOverlap 0.8’. For spike-in controlled analysis, reads mapped to drosophila genes were also counted using the same method. Differential gene expression analysis was performed using DESeq2 (*57*). Spike-in normalization was incorporated by calculating size factors using ‘estimateSizeFactors’ function based on drosophila gene read counts. Human genes with fold-change (FC) > 1.5 and false discovery rate (FDR) < 0.05 were selected as differentially expressed genes for downstream analysis. Gene ontology (GO) analysis was done using EnrichR (*58*), where top significant GO terms by adjusted p-value were selected for presentation.

### CUT&RUN, ChIP-seq, and ATAC-seq data analysis

CUT&RUN Sequencing reads were aligned to the same genome reference as RNA-seq data for consistency using STAR aligner like RNA-seq with options ‘--alignSJDBoverhangMin 999 -- alignIntronMax 1 --alignMatesGapMax 2000 --outFilterMultimapNmax 1 -- outFilterMismatchNoverLmax 0.05’. Only uniquely and concordantly aligned read pairs were used for analysis. Read 1 and 2 were connected with each other to form a fragment. Downstream analysis was performed largely as previously described (*59*). For transcription factor (TF) CUT&RUN, fragments with size < 120 bp were selected as originated from putative nucleosome-free regions. For histone modification (histone) CUT&RUN, fragments with size > 150bp were selected as originating from putative nucleosomal regions. Selected fragments were resized to 100bp to normalize bias from the sample-to-sample variabilities of fragment lengths. Peak calling was done using Homer (*60*) against a matching IgG control. Target-specific options were used for peak calling: ‘-style factor -center -size 200 -tbp 0 -fragLength 100’ for TF and ‘-style histone - tbp 0 -fragLength 100’ for histone. Peaks overlapping with ENCODE blacklist regions (*61*) were discarded. For TF, peaks with > 1rpm were retained for downstream analysis. Bigwig files were generated using bedtools (*62*) and UCSC toolkit (*63*) in RPM scale.

ChIP-seq sequencing reads were aligned to the same genome reference using STAR aligner with options ‘--alignSJDBoverhangMin 999 --alignIntronMax 1 --alignMatesGapMax 2000 -- outFilterMultimapNmax 1 --outFilterMismatchNoverLmax 0.05’. Redundant fragments were deduplicated using Picard (https://broadinstitute.github.io/picard). BigWig files were generated similarly but without fragment length restriction. PRDM Peak calling was done using Homer against matching an input sample, and peaks > 0.7RPM were retained for downstream analysis.

ATAC-seq data from endoderm development were downloaded from GEO (GSE114101) and processed in a similar way with CUT&RUN to identify open chromatin peaks and to create bigwig files for visualization. ChIP-seq data from the public domain was downloaded from GEO. Data were either processed from scratch following the standard pipeline (STAR aligner and Homer peak calling) or converted to hg38 using liftOver (*63*) if processed data is available only in hg19.

Pioneer and PRDM factors were defined as “co-bound” if the distance between the peak centers is < 2kb. Putative target genes of TF-bound regions were defined by GREAT analysis (*21*). Genomic annotations of TF peaks were performed using Homer. All de novo motif searches were done using Homer within 200bp window.

Line plot visualization of CUT&RUN and ChIP-seq data was done by extracting RPM-normalized stack-height profile from bigwig files using bwtool (*64*) and by drawing using customized R scripts. ChIP-seq data were input-subtracted before statistical testing or visualization. Wilcoxon tests in the boxplots were performed by taking the average meta-profile within the corresponding window per genomic loci and comparing them between groups.

**Fig. S1.**
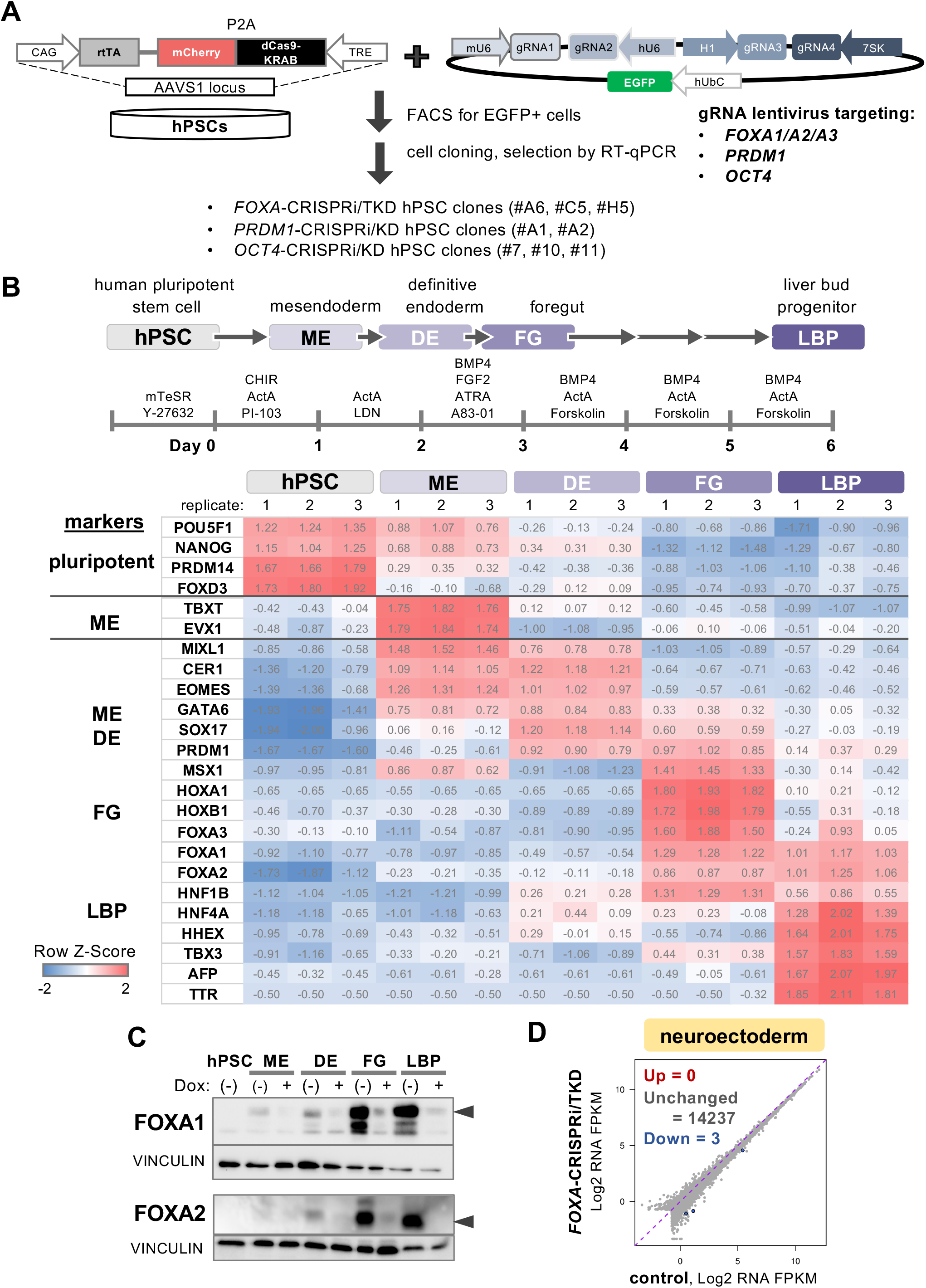
Establishment of doxycycline (dox) inducible FOXA-CRISPRi/TKD knockdown system in human endoderm differentiation. **(A)** Schematic overview of of the strategy for generating dox-inducible CRISPRi hPSCs. The cassette containing dox-controlled reverse transcriptional activator (rtTA) driven by a CAG promoter and the dCas9-KRAB-P2A-mCherry driven by a dox/tet-response element (TRE) was integrated into AAVS1 locus of hPSCs. gRNAs targeting *FOXA1/A2/A3, PRDM1*, or *OCT4* gene locus were cloned into a lentivirus vector containing EGFP reporter. **(B)** Schematic of stepwise endoderm differentiation protocol from hPSCs. RNA-seq heatmap of representative marker gene expression during the endoderm differentiation in Z-score. **(C)** Western blot analysis of FOXA1 and FOXA2 proteins along with internal control, VINCULIN protein, in control and *FOXA*-TKD cells. **(D)** Differential gene expression analysis of RNA-seq comparing *FOXA*-TKD neuroectoderm versus control (n=2 replicates from 2 independent *FOXA*-CRISPRi clones; FDR < 0.05, FC > 1.5).

**Fig. S2.**
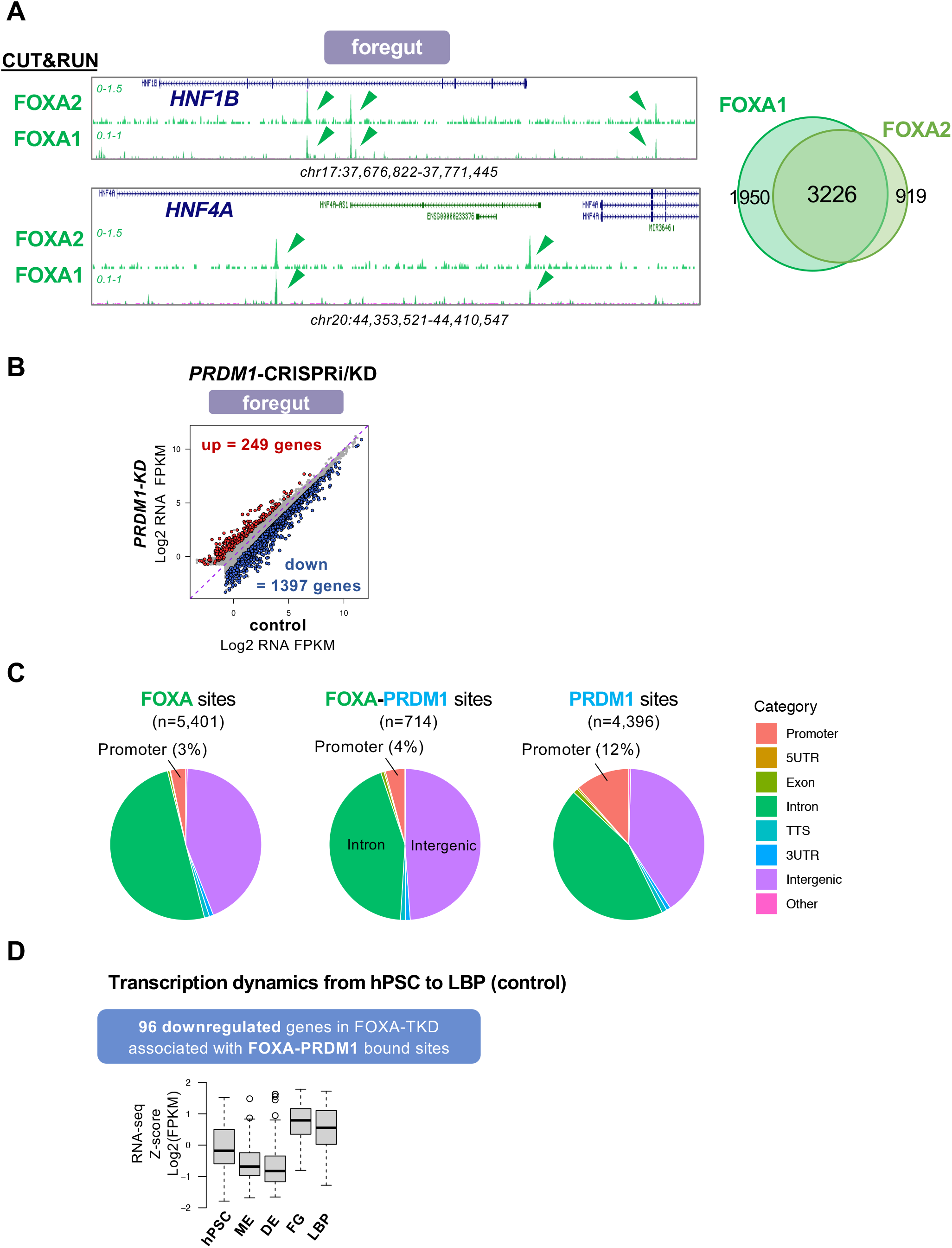
Characterization of FOXA and PRDM1 binding sites and their target genes. **(A)** FOXA1 and FOXA2 CUT&RUN tracks in control foregut cells at *HNF1B* and *HNF4A* loci in RPM scale. **(B)** Differential gene expression analysis of RNA-seq comparing *PRDM1*-CRISPRi/KD foregut cells versus control (n=2 replicates from 2 independent *PRDM1*-CRISPRi clones; FDR < 0.05, FC > 1.5). **(C)** Genomic annotations of the FOXA-only bound sites, FOXA-PRDM1 bound sites, and PRDM1-only bound sites by Homer. TTS, transcription termination site. **(D)** Transcriptional dynamics of FOXA-PRDM1 targeting genes during normal endoderm differentiation.

**Fig. S3.**
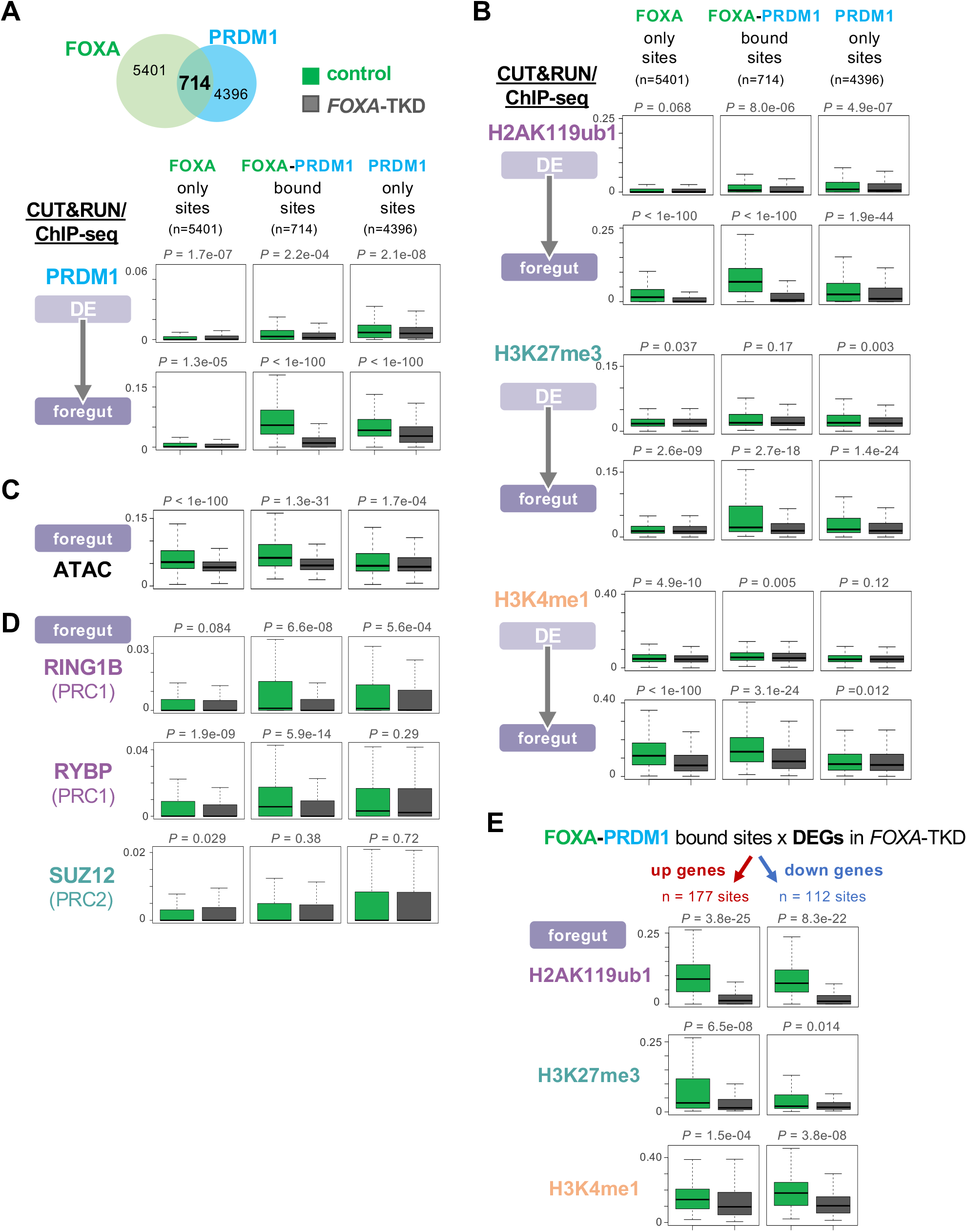
Characterization of FOXA and PRDM1 bound enhancers. **(A** and **B)** Changes of PRDM1 ChIP-seq, H2Aub ChIP-seq, H3K27me3 CUT&RUN, and H3K4me1 CUT&RUN signals in control (green boxplots) and *FOXA-*TKD (gray boxplots) definitive endoderm (DE) and foregut at FOXA-only bound sites, FOXA-PRDM1 bound sites, and PRDM1-only bound sites. **(C)** Changes of ATAC-seq signal in control (green boxplots) and *FOXA2* knockout (gray boxplots) foregut cells. **(D)** Changes of RING1B, RYBP, and SUZ12 ChIP-seq signals in control (green boxplots) and *FOXA-*TKD (gray boxplots) foregut cells. **(E)** Changes of H2Aub, H3K27me3, and H3K4me1 signals in control (green boxplots) and *FOXA*-TKD (gray boxplots) foregut at FOXA-PRDM1 bound sites associated with differentially regulated genes in *FOXA*-TKD. *P* value by Wilcoxon test.

**Fig. S4.**
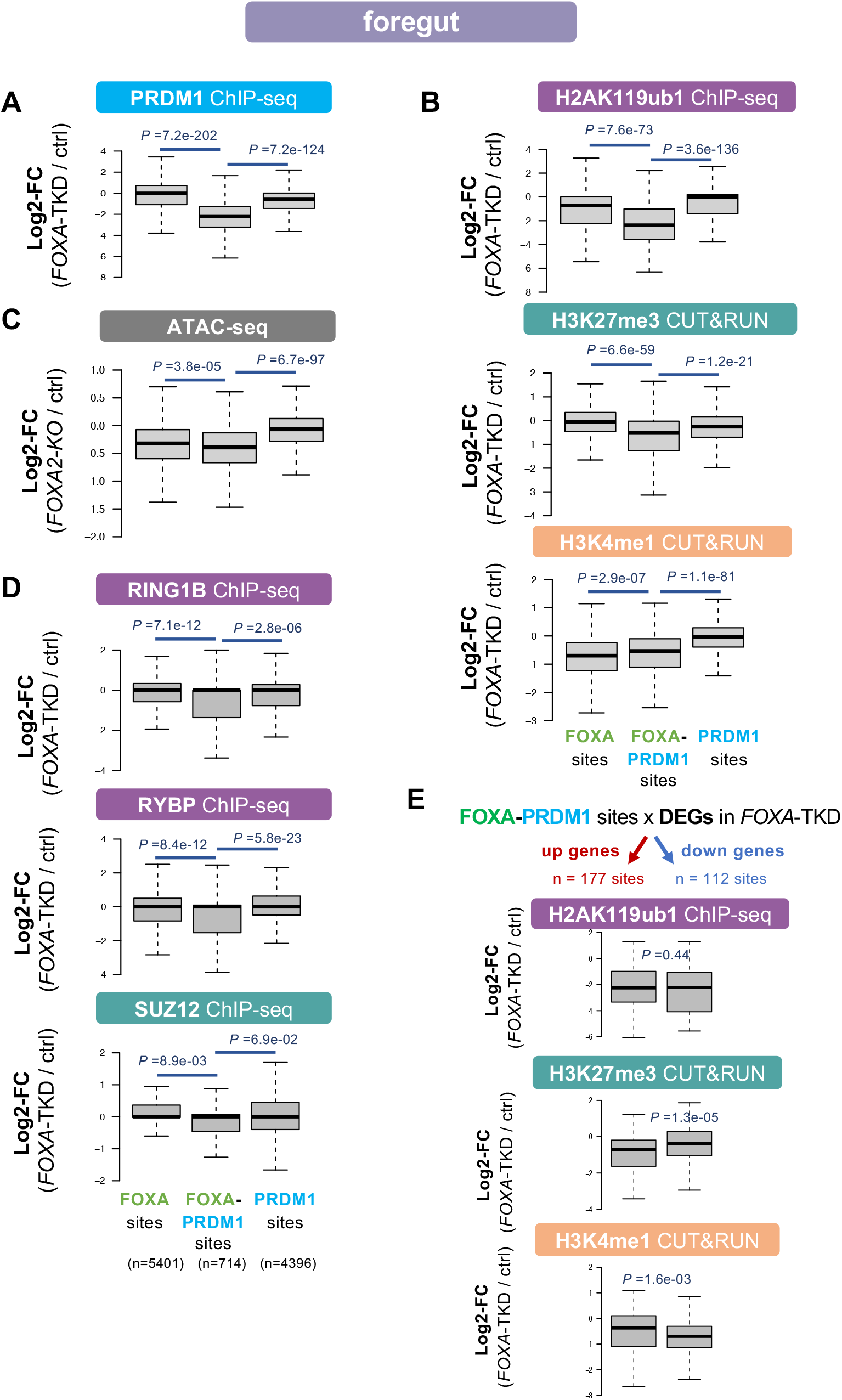
Characterization of FOXA and PRDM1 bound enhancers. **(A** and **B)** Log2 fold change (FC) of PRDM1 ChIP-seq, H2Aub ChIP-seq, H3K27me3 CUT&RUN, and H3K4me1 CUT&RUN signals in *FOXA*-TKD against control foregut cells at FOXA-only bound sites, FOXA-PRDM1 bound sites, and PRDM1-only bound sites. **(C)** Log2 fold change of ATAC-seq signal in *FOXA2* knockout against control foregut cells. **(D)** Log2 fold change of RING1B, RYBP, and SUZ12 ChIP-seq signals in *FOXA*-TKD against control foregut cells. **(E)** Log2 fold change of H2Aub, H3K27me3, and H3K4me1 signals in *FOXA*-TKD against control foregut cells at FOXA-PRDM1-both bound sites associated with differentially regulated genes in *FOXA*-TKD. *P* value by Wilcoxon test.

**Fig. S5.**
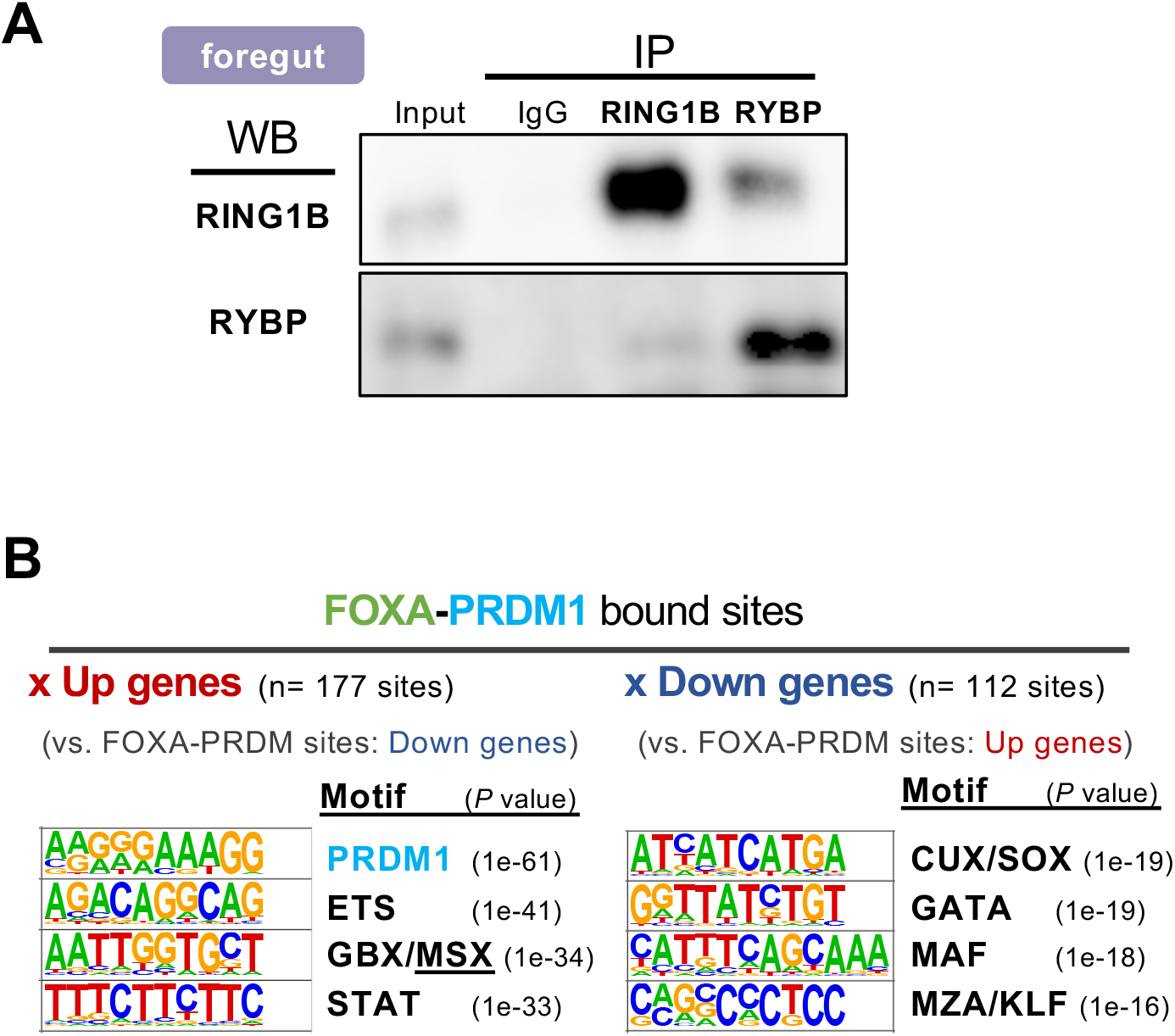
Characterization of FOXA and PRDM1 bound enhancers. **(A)** Western blot (WB) validation of successful co-immunoprecipitation (IP) reaction for RING1B and RYBP. **(B)** Top motifs enriched at FOXA-PRDM1 bound peaks associated with upregulated genes against a background of FOXA-PRDM1 peaks associated with downregulated genes, and vice-versa, by HOMER.

**Fig. S6.**
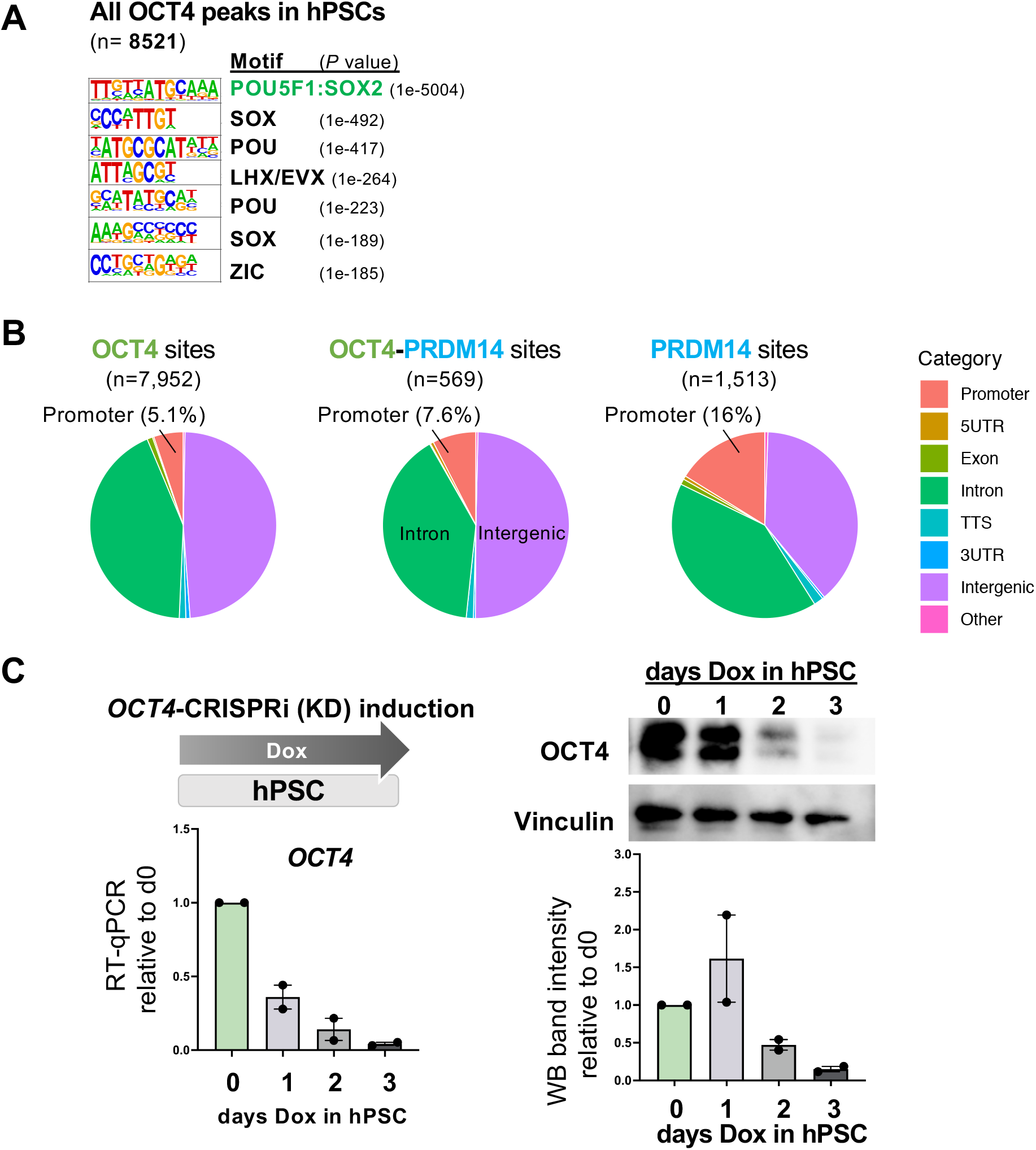
Characterization of OCT4 and PRDM14 binding sites and validation of *OCT4*-CRISPRi/KD hPSCs. **(A)** Top motifs enriched at OCT4 ChIP-seq peaks in control hPSCs by HOMER. **(B)** Genomic annotations of the OCT4-only bound sites, OCT4-PRDM14 bound sites, and PRDM14-only bound sites by Homer. TTS, transcription termination sites. **(C)** Validation of *OCT4*-CRISPRi/KD knockdown efficiency by RT-qPCR and Western blot (WB) assays in hPSCs (n=2 replicates from 2 independent CRISPRi clones, Means ± SD).

**Fig. S7.**
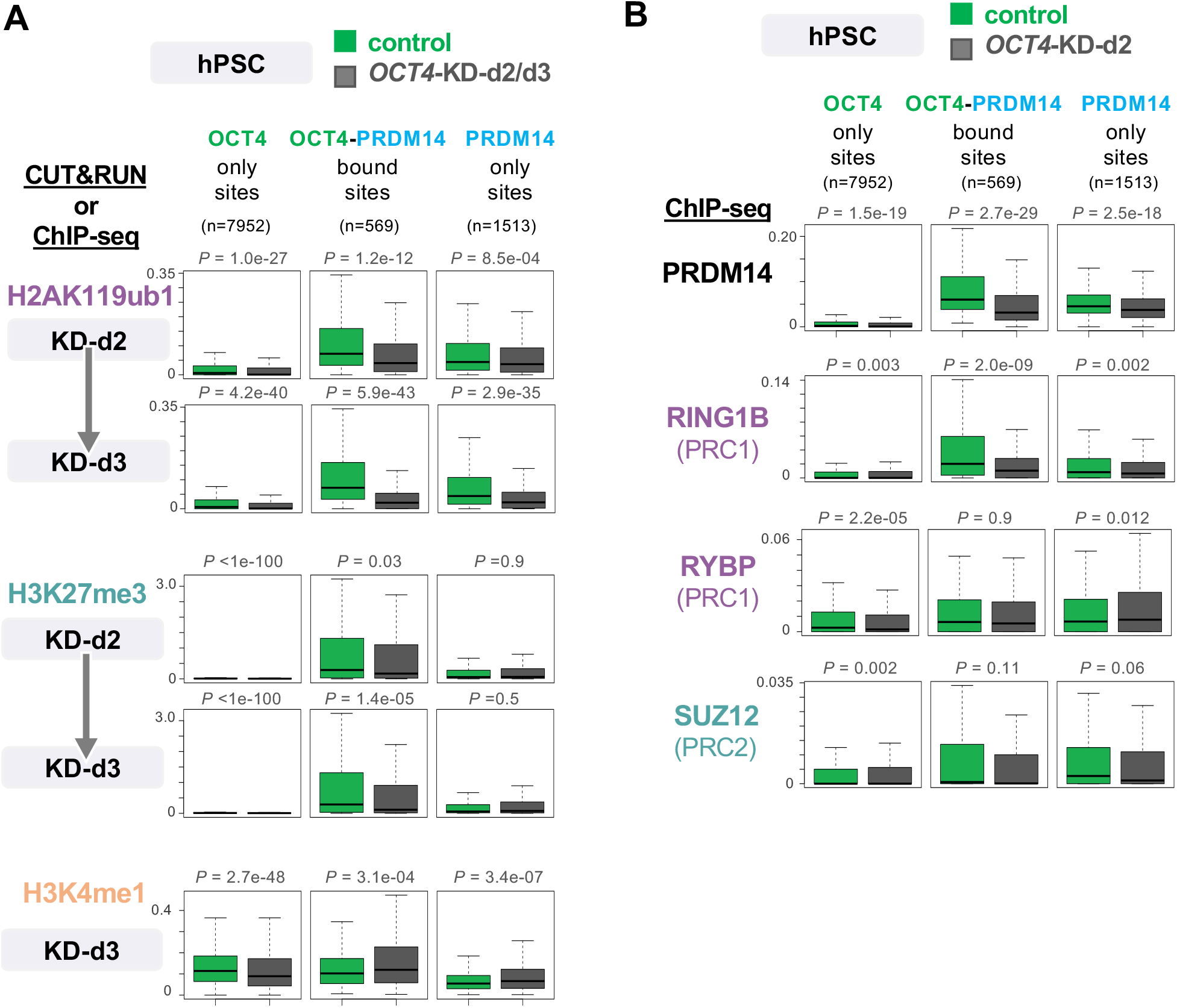
Characterization of OCT4 and PRDM14 bound enhancers. **(A)** Changes of H2Aub ChIP-seq, H3K27me3 CUT&RUN, and H3K4me1 CUT&RUN signals in control (green boxplots) and *OCT4-*KD-d2/d3 (gray boxplots) hPSCs at OCT4-only bound sites, OCT4-PRDM14 bound sites, and PRDM14-only bound sites. **(B)** Changes of PRDM14, RING1B, RYBP, and SUZ12 ChIP-seq signals in control (green boxplots) and *OCT4-*KD-d2 (gray boxplots) hPSCs. *P* value by Wilcoxon test.

**Fig. S8.**
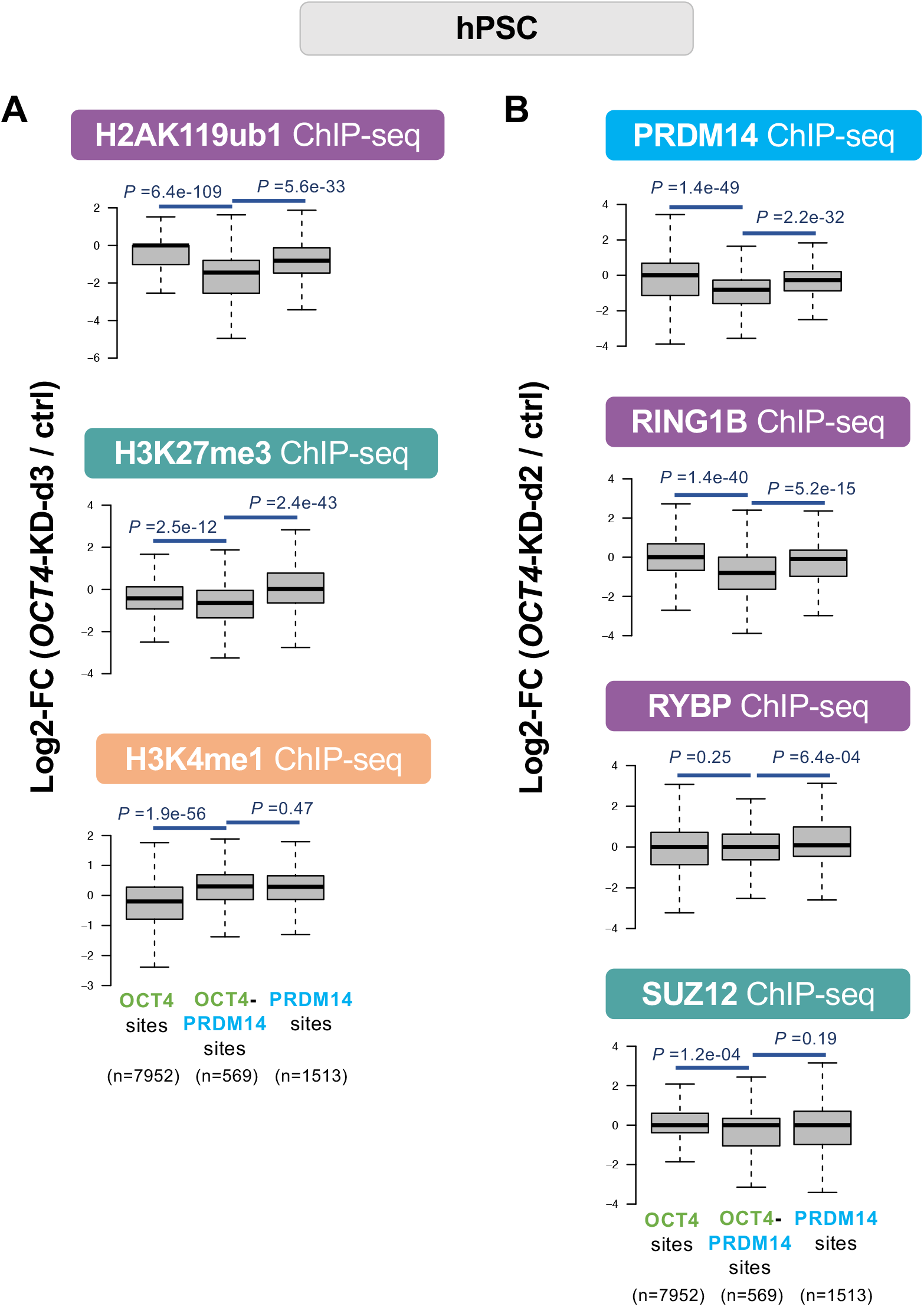
Characterization of OCT4 and PRDM14 bound enhancers. **(A)** Log2 fold change (FC) of H2Aub ChIP-seq, H3K27me3 CUT&RUN, and H3K4me1 CUT&RUN signals in *OCT4-*KD-d3 against control hPSCs at OCT4-only bound sites, OCT4-PRDM14 bound sites, and PRDM14-only bound sites. **(B)** Log2 fold change of PRDM14, RING1B, RYBP, and SUZ12 ChIP-seq signals in *OCT4-*KD-d2 against control hPSCs. *P* value by Wilcoxon test.

